# Diversification of CORVET tethers facilitates transport complexity in *Tetrahymena thermophila*

**DOI:** 10.1101/737247

**Authors:** Daniela Sparvoli, Martin Zoltner, Mark C. Field, Aaron Turkewitz

**Author notes:** (MZ) Biotechnology and Biomedicine Centre of the Academy of Sciences and Charles University (BIOCEV), Průmyslová 595, 252 50 Vestec, Czech Republic.

## Abstract

In endolysosomal networks, two hetero-hexameric tethers called HOPS and CORVET are found widely throughout eukaryotes. The unicellular ciliate *Tetrahymena thermophila* possesses elaborate endolysosomal structures, but curiously both it and related protozoa lack the HOPS tether and several other trafficking genes while retaining the related CORVET complex. *Tetrahymena* encodes multiple paralogs of most CORVET subunits, which assemble into six distinct complexes. Each complex has a unique subunit composition and, significantly, shows unique localization, indicating participation in distinct pathways. One pair of complexes differ by a single subunit (Vps8), but have late endosomal vs. recycling endosome locations. While Vps8 subunits are thus prime determinants for targeting and functional specificity, determinants exist on all subunits except Vps11. This unprecedented expansion and diversification of CORVET provides a potent example of tether flexibility, and illustrates how ‘backfilling’ following secondary losses of trafficking genes can provide a mechanism for evolution of new pathways.

## Background

Nearly all eukaryotic cells contain multiple membrane-bound compartments, with either an endogenous or endosymbiotic origin. Bidirectional transport of proteins and lipids between compartments is critical for cell function and defects contribute towards pathologic disorders including neurodegeneration and cancer(Mellman and Yarden, 2013; Neefjes and van der Kant, 2014). The endolysosomal network refers to a subset of pathways linking endocytic traffic with degradative and secretory compartments(Huotari and Helenius, 2011). To ensure accurate trafficking cells deploy intricate mechanisms to license productive interactions between compartments that ultimately allow content mixing *via* membrane fusion(Kummel and Ungermann, 2014).

A key protein determinant for ensuring faithful membrane interactions are transmembrane SNARE proteins, with distinct paralogs present at each compartment(Gerst, 1999). SNARE complex assembly drives formation of a fusion pore, together with complexes called tethers that act upstream of SNAREs(Baker and Hughson, 2016). The HOPS (homotypic fusion and protein sorting) and CORVET (class C core vacuole/endosome tethering) tether complexes, first discovered in *S. cerevisiae*, are cytoplasmic hetero-hexamers that bridge compartments by binding Rab GTPases at both transport vesicle and destination compartment membranes, and subsequently chaperoning SNARE assembly(Baker and Hughson, 2016; Horazdovsky et al., 1996; Nickerson et al., 2009; Spang, 2016; van der Beek et al., 2019). These reactions have been reconstituted in elegant *in vitro* systems, but comparably detailed *in vivo* characterization remains lacking, particularly for CORVET(Ho and Stroupe, 2016; Lobingier and Merz, 2012; Lobingier et al., 2014; Orr et al., 2017; Schwartz et al., 2017). CORVET and HOPS mediate endosome maturation through interaction with Rab5/Vps21 and Rab7/Ypt7 respectively, promoting fusion of early endosomes (EE) with multivesicular late endosomes (LE) and subsequently vacuoles/lysosomes (LL)(Balderhaar and Ungermann, 2013). HOPS and CORVET share four core subunits: Vps11 (Vacuolar Protein Sorting), Vps16, Vps18 and Vps33(Nickerson et al., 2009). In addition, each complex contains two specific subunits: Vps3 and Vps8 for CORVET and Vps39 and Vps41 in HOPS(Peplowska et al., 2007). One current model is that CORVET may convert to HOPS during endosome maturation by exchanging the complex-specific subunits(Ostrowicz et al., 2010; Peplowska et al., 2007). It is particularly compelling since the complex-specific subunits directly bind Rabs, and thus are key to CORVET vs HOPS specificity(Markgraf et al., 2009). Intriguingly, hybrid CORVET/HOPS complexes have been identified in *S. cerevisiae*(Ostrowicz et al., 2010; Peplowska et al., 2007), but these are low abundance and only detected under over-expression conditions, and hence may represent the product of high concentration driving low affinity interactions.

While budding yeast has single genes encoding each CORVET and HOPS subunit, the genetic and cell biological landscapes have additional dimensions in metazoa. Two paralogs of VPS33 are present in mammals, zebrafish, *Drosophila,* and *C. elegans* (Gissen et al., 2005); similarly, two VPS16 paralogs are present in mammals and flies. In *C. elegans*, the two Vps33 paralogs belong to HOPS and CORVET, respectively (Solinger and Spang, 2014), while in mammals and flies the pairs of Vps33 and Vps16 paralogs belong to alternative complexes. In mammals, Vps16A and Vps33A belong to HOPS and CORVET, while Vps16B (also called VIPAR/VIPAS39) and Vps33B are part of a related but functionally distinct complex called CHEVI (class C Homologues in Endosome–Vesicle Interaction) (Spang, 2016). CHEVI is associated with both conserved, lineage- and tissue-specific structures, e.g., α-granules and lamellar bodies, which are mammalian platelet-specific and keratinocyte-specific lysosome-related organelles (LROs)(Bem et al., 2015; Dai et al., 2016; Lo et al., 2005; Rogerson and Gissen, 2018). A similar configuration pertains for *Drosophila* Vps16B and Vps33B, which also form a novel complex (Cullinane et al., 2010; Gissen et al., 2004; Hunter et al., 2018; Pulipparacharuvil et al., 2005; Tornieri et al., 2013). Further, HOPS or CORVET subunits may function in stable sub-complexes. For example, a Vps3-Vps8 subcomplex in Hela cells serves a CORVET-independent function in recycling internalized β1-integrins from early to recycling endosomes(Jonker et al., 2018). In *Drosophila*, miniCORVET is comprised of Vps8, Vps16, Vps18 and Vps33, and functions at early endosomes (Lorincz et al., 2016), while mammalian Vps41, in the absence of other HOPS subunits, is involved in sorting cargo to the regulated secretory pathway (Asensio et al., 2013).

Taken together, these studies reveal remarkable flexibility in HOPS/CORVET subunits contributing to a wide range of functions. Nonetheless, virtually all detailed studies have been pursued in a single eukaryotic lineage, the Opisthokonts, which includes both fungi and all of the animals discussed above(Lynch et al., 2014). Hence, the potential full diversity of HOPS and CORVET structure and function remains unexplored. Interestingly, recent studies in the Archaeplastida (plants) reveals that the coupling between CORVET and HOPS functions may itself be evolutionarily plastic(Takemoto et al., 2018).

Ciliates are very distantly related to either Opisthokonts or Archaeplastids. Together with Dinoflagellates and Apicomplexans, ciliates constitute the Alveolate branch of the Stramenopile–Alveolate–Rhizaria (SAR) supergroup(Adl et al., 2012). Many of these organisms are free-swimming protozoa whose estimated 30,000 morphologically-diverse species(Adl et al., 2007) play key ecological roles in a wide range of freshwater, marine and terrestrial environments (Gimmler et al., 2016; Warren et al., 2017; Weisse, 2017; Zingel et al., 2019). Ciliates also exhibit striking cellular and behavioral complexity for single-celled organisms, and ciliate genomes encode correspondingly large numbers of genes(Hausmann, 1996; Wang et al., 2017). For example, *Tetrahymena thermophila* expresses roughly the same number of Rab GTPases as in humans, hinting at the elaboration of membrane trafficking pathways(Bright et al., 2010; Saito-Nakano et al., 2010). This sophistication in *Tetrahymena* is also manifest in capacities also present in mammalian cells, such as the ability to store peptides and release them in a stimulus-dependent manner(Guerrier et al., 2017).

Although molecular studies are scant, ciliates maintain an elaborate endolysosomal network. Morphological studies suggest at least four distinct pathways for uptake in *Tetrahymena* including clathrin-mediated endocytosis(Elde et al., 2005; Nilsson and Van Deurs, 1983) and phagosome formation at an anterior portal called the oral apparatus, followed by phagolysosome maturation *via* fusion with multiple classes of endosomes (Jacobs et al., 2006; Nilsson, 1979; Plattner, 2010). Other less well-characterized modes of uptake are associated with recovery following exocytic events(Frankel, 2000). Ciliates also have autophagy-related phenomena that depend on endolysosomal trafficking, particularly prominent during the stage in conjugation when selected nuclei are eliminated (Akematsu et al., 2014; Davis et al., 1992; Liu and Yao, 2012; Orias et al., 2011). Another relevant structure is the contractile vacuole, a water-pumping organelle (Allen, 2000; Plattner, 2015) that, based on its associated Rab proteins, likely belongs to the endolysosomal network (Bright et al., 2010). Lastly, prominent secretory vesicles in *Tetrahymena* called mucocysts are LROs(Briguglio et al., 2013; Kaur et al., 2017).

*Tetrahymena* and related ciliates possess an atypical complement of genes encoding endolysosomal tethers and other trafficking genes, indicating a significant bottleneck in ancestors of this lineage that resulted in secondary losses, and particularly in Rab GTPases(Sparvoli et al., 2018). Most relevant here is that both HOPS-specific subunits were lost, but most other subunits (i.e., the core subunits, as well as the CORVET-specific subunits) were retained and expanded into multiple paralogs(Klinger et al., 2013). *T. thermophila* expresses two Vps33, two Vps16, two Vps3, four Vps18 and six Vps8 paralogs. We previously discovered, starting from a forward genetic approach, that the “a” paralog of Vps8 was essential for mucocyst formation(Sparvoli et al., 2018). In particular, *Δvps8a* cells failed to deliver a subset of mucocyst cargo proteins, consistent with the idea that a specific pathway depended upon the activity of a Vps8a-containing tether. Taken together with genomic data, these results suggested that a large evolutionary expansion of pathway-specific CORVET-related tethering complexes in a family of ciliates accompanied elaboration of a complex endolysosomal network, following a reduction in complexity. To encapsulate this idea, we suggest the term ‘backfilling’, whereby the retained components expand to occupy a gap in functionality arising from gene loss.

Our previous data did not directly address whether Vps8a belonged to the inferred CORVET complex and if so, whether the six Vps8 paralogs define biochemically or functionally distinct complexes. Here we provide evidence that *Tetrahymena* assembles six distinct CORVET complexes, and that each hexameric complex consists of a unique combination of subunit paralogs. The only subunit for which a single protein contributes to all complexes is Vps11, which is a key integrator of HOPS and CORVET assembly in other organisms(Plemel et al., 2011). The six *Tetrahymena* CORVET complexes function at six distinct cellular locations including phagosomes at two different maturation stages, and contractile vacuoles. Steady-state localization of complexes is determined by Vps8, since complexes differing only in that subunit show unique localization. Analysis of the subunit composition of the cohort further indicates that Vps18 plays a key role in shaping the specificity of complex assembly.

## Results

### Vps8ap defines a specific hexameric CORVET complex in *Tetrahymena*

To ask whether Vps8a is associated with additional subunits forming a classical CORVET complex, we characterized Vps8a-associated proteins using biochemical and physical approaches. We first asked whether Vps8a associated with other canonical CORVET subunits by co-expressing FLAG-tagged Vps8a with 6myc-tagged versions of several core subunits. We then precipitated Vps8a and evaluated co-precipitation of the 6myc-tagged proteins. To avoid possible non-specific interactions due to overexpression, we tagged all subunits by integrating epitope tags at the 3’ ends of their endogenous loci.

Prior annotation of CORVET subunits encoded in the *T. thermophila* genome revealed multiple paralogs for most subunits including two for *VPS16*, four *VPS18* and two *VPS33*(Klinger et al., 2013). Single genes were reported for *VPS11* and *VPS3*, although a second *VPS3* gene is in fact present (see below). To begin our analysis, we asked whether Vps8a associated with Vps16, 18, or 33. We tagged all paralogs for each subunit in pairwise combination with Vps8a-FLAG, to ask whether the paralogs were differentially associated with Vps8a. The results showed that Vps16b, Vps33b, and Vps18d, but not the other paralogs of each subunit, could be robustly co-precipitated with Vps8a (Fig. 1A).

**Figure 1.**
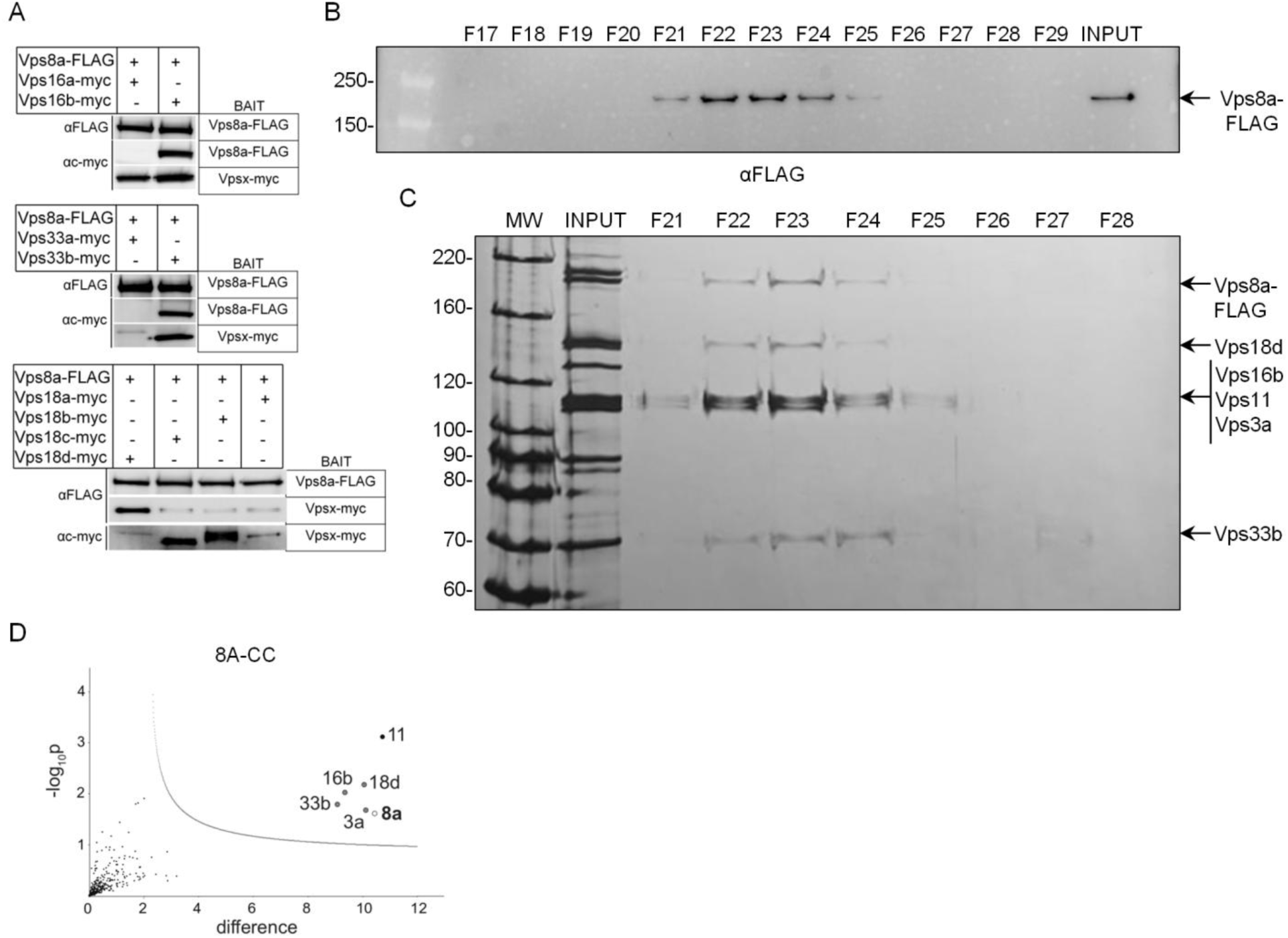
Vp8a associates with five other subunits in a hexameric CORVET complex (8A-CC). A) Co-immunoprecipitation of Vps8a-FLAG with myc-tagged Vp16b, Vp18d, and Vp33b subunits. Cells were transformed to endogenously express Vps8a-FLAG in pairwise combination with myc-tagged Vps16a or b, Vps33a or b, and Vps18a, b, c or d. Detergent cell lysates were split, to be incubated with anti-c-myc beads or anti-FLAG beads. Samples were analyzed by SDS-PAGE and Western-blotted with mAb anti-c-myc and anti-FLAG antibodies. B) and C) Sedimentation analysis of 8A-CC. B) Vps8a-FLAG and associated proteins were immunoisolated using anti-FLAG beads, from suspended cryopowders derived from cells expressing Vps8a-FLAG. Eluted proteins were applied to 11-ml 10%-40% continuous glycerol gradients, and centrifuged at 235,000 x g for 18h at 4°C. 250µl fractions were collected, of which 25µl aliquots were subjected to SDS-PAGE (4-20% gel). Fractions 17-29 are shown. Vp8a was detected by Western blot with anti-FLAG antibodies. “INPUT” corresponds to 1% of the total eluate. C) 4% of the total eluate (INPUT) and 35µl aliquots of the gradient fractions (F21 to F28) were subjected to SDS-PAGE (8% gel) and visualized by silver staining. The Vps8a subunit, as well as bands of the expected sizes for five additional CORVET subunits, are identified at the right. Thyroglobulin, run in parallel as a size standard, appeared in fractions 24-29, with a peak in fraction F26. D) Mass spectrometric identification of proteins co-isolated with Vps8a. Cryopowders (150g) from wild-type and Vps8a-FLAG expressing cells were solubilized and treated as in (B), except bound proteins were eluted with LDS-sample buffer. The total eluates were applied to 4-20% gels for SDS-PAGE and allowed to migrate ∼1cm, after which a single 1 cm gel slice per lane was processed for mass-spectrometric analysis. On Volcano plots such as the one shown here, proteins falling above the threshold line are considered significant. The open circle marks the Vps8a subunit used as bait, while the black circle marks the unique Vps11 subunit. The light grey circles indicate specific paralogs of the other four CORVET subunits. Each sample was prepared in duplicate.

Our results suggest that Vps8a belongs to a canonical hexameric complex, but smaller sub-complexes may also exist as described above. To examine this possibility, we immunoisolated the Vps8a containing CORVET complex by affinity capture using Vps8a-FLAG as bait. To overcome any issues from low expression levels we adapted methods used in trypanosomes (Obado et al., 2016), yeast (Oeffinger et al., 2007) and mammalian cells(LaCava et al., 2016), where large numbers of cells are rapidly frozen and milled to generate cryopowders. We tested a variety of buffer conditions for solubilizing Vps8a from the cryopowders and final conditions resulted in ∼60% solubilization (not shown). Vps8a and associated proteins were immunoisolated from cryopowder solutes and eluted with excess FLAG peptide, followed by glycerol gradient centrifugation. We identified the fractions containing Vps8a by western blotting (Fig. 1B), and visualized the co-sedimenting proteins in those fractions by silver staining (Fig. 1C). Notably peak fractions 22 and 23 contained prominent bands at the expected sizes for Vps16b, Vps33b, and Vps18d, and also for Vps11 and Vps3a, suggesting that a full hexameric complex was present. However, as Vps16b, Vps3a and Vps11 are all between 100-120kDa, they are difficult to resolve (Fig.1C). Importantly, no Vps8a was detected in lower fractions (not shown), suggesting that this subunit is largely associated with a stable hexameric complex.

To confirm our interpretation of the glycerol gradient centrifugation and to identify the specific gene products, we immunoisolated Vps8a and associated proteins and analyzed the eluate by liquid chromatography-tandem mass spectrometry (LC-MS/MS). Vps8a-associated proteins prominently include Vps16b, Vps33b, Vps18d, Vps3a and Vps11, with no other paralogs detected (Figure 1D). Taken together, our results indicate the Vps8a belongs to a hexameric CORVET complex, which we refer to as 8A-CC. However, we note that the non-soluble fraction of Vps8a could potentially participate in a different biochemical complex.

### Vps8b-f associate with distinct subunit combinations and *Tetrahymena* expresses six distinct CORVET complexes

Vps8a is one of six Vps8 paralogs expressed in *Tetrahymena*. These paralogs are ancient in origin: the split between even the most-closely related pair (*VPS8A* and *VPS8C*) predates the ∼22MYA divergence between *T. thermophila* and *T. malaccensis*(Sparvoli et al., 2018). Moreover, these paralogs have been maintained in multiple species suggesting that they provide important, non-redundant functions. The six *VPS8* paralogs in *T. thermophila* differ in their transcriptional profiles, consistent with functional diversification(Sparvoli et al., 2018). To determine if each Vps8 paralog belongs to a unique biochemical complex, we expressed each as an endogenously-FLAG-tagged fusion. The full length constructs were confirmed by immunoprecipitation, followed by SDS-PAGE and western blotting (Fig. S1A). We then used cryomilling and immunoisolation as above, followed by SDS-PAGE. In silver-stained polyacrylamide gels of the eluted complexes, we detected multiple bands in the size range expected for CORVET subunits (Fig. S1B), but many bands had distinct migrations from the 8A-CC pulldown bands (Fig. 1C).

We analyzed each of these immunoisolated mixtures by LC-MS/MS (Figure 2A). For each Vps8 paralog, the most enriched proteins consisted of five canonical CORVET subunits, as expected for hexameric complexes. In all cases, a single paralog for each subunit was identified, echoing our findings for 8A-CC. Based on this, the composition of these complexes, which we call 8B-CC, 8C-CC, etc., can be predicted with confidence (Fig. 2B, Table 1). As a cohort, the CORVET complexes share just one gene product, encoded by *VPS11*. In yeast CORVET and HOPS complexes, Vps11 has a key role in complex assembly(Ostrowicz et al., 2010; Plemel et al., 2011). Furthermore, informatics-based analysis suggested that all CORVET complexes contained the identical Vps3 subunit, as only one VPS3 gene was identified(Klinger et al., 2013). However, the LC-MS/MS revealed that 8B-CC contains a distinct and divergent Vps3 paralog.

**Figure 2.**
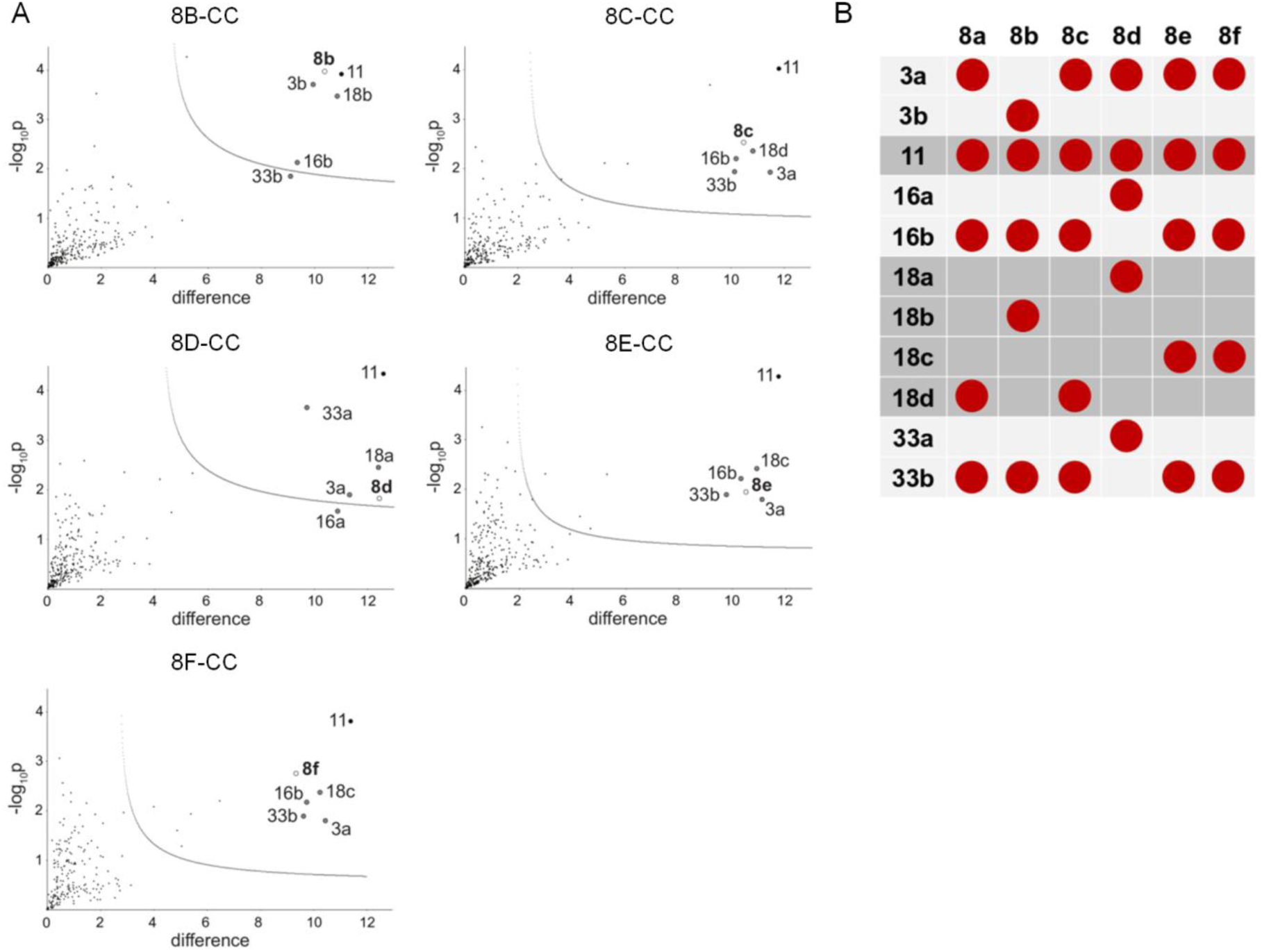
*Tetrahymena* has six unique CORVET complexes. A) Volcano plots of mass spectrometry results, identifying the CORVET subunit paralogs associated with FLAG-tagged Vps8b, 8c, 8d, 8e, and 8f. All samples were prepared as in Figure 1D, in duplicate. Significant hits are shown above the threshold line in each plot. B) Dotplot showing the comprehensive composition of CORVET complexes in *T. thermophila*. Each of the six Vps8 paralogs (top row) is associated with five other subunits, whose identities are indicated in the left column. 8A-CC and 8C-CC share five subunits, as do 8E-CC and 8F-CC. 8B-CC possesses unique Vps8, Vps3, and Vps18 subunit paralogs. 8D-CC possesses unique paralogs of the Vps8, 16, 18, and 33 subunits.

**Table 1.** Gene IDs of CORVET subunits

| TTHERM_ID | Gene Name |
| --- | --- |
| 00290710 | *VPS8A* |
| 00393150 | *VPS8B* |
| 00716100 | *VPS8C* |
| 00532700 | *VPS8D* |
| 00691590 | *VPS8E* |
| 00384890 | *VPS8F* |
| 01141590 | *VPS3A* |
| 000777289 | *VPS3B* |
| 01205260 | *VPS11* |
| 00370870 | *VPS16A* |
| 00294740 | *VPS16B* |
| 00242090 | *VPS18A* |
| 00463740 | *VPS18B* |
| 00292270 | *VPS18C* |
| 00410250 | *VPS18D* |
| 00355020 | *VPS33A* |
| 000077749 | *VPS33B* |

Two pairs of complexes are closely related in subunit composition; 8A- and 8C-CC are identical except for the Vps8, with the same relationship existing between 8E- and 8F-CC. By contrast 8B-CC and 8D-CC each show an exclusive combination of subunits. Interestingly, the composition of the six CORVET complexes appears largely consistent with evolutionary relationships previously inferred(Sparvoli et al., 2018), in that more closely related Vps8 paralogs can now be seen to belong to complexes that share a larger number of identical subunit paralogs.

### The six *Tetrahymena* Vps8 paralogs have distinct locations

To understand how six biochemically-distinct CORVET tethers contribute to cellular function, we first asked whether they associate with different compartments. We generated cell lines where mNeon was integrated immediately downstream of each *VPS8* open reading frame, to express endogenous levels of tagged protein. These integrated constructs were all driven to fixation, to completely replace the wildtype alleles in the somatic macronuclei. Fusion proteins of the expected sizes were detected by immunoprecipitation followed by SDS-PAGE and western blotting, though proteolytic cleavage of some products was also seen, discussed further below (Fig. S2A). *VPS8C* and *D* are essential genes(Sparvoli et al., 2018). Since the cells relying on mNeon-tagged Vps8c and 8d had no detectible growth phenotypes, we infer that tagging does not interfere with protein activity. Consistent with this, Vps8a-mNeon is fully functional(Sparvoli et al., 2018).

We analyzed the localization of mNeon-tagged Vps8 paralogs under a variety of conditions. First, cells were transferred for 2h to a medium that reduces the autofluorescence in food vacuoles, and then immobilized in agarose dissolved in a Tris buffer (Fig. 3A). In other experiments, cells in standard growth medium were fixed for imaging (Fig. 3B). All six Vps8 paralogs are expressed at low levels (TetraFGD; http://tfgd.ihb.ac.cn)(Xiong et al., 2011a; Xiong et al., 2013), so that detecting the mNeon-fusions in whole cell lysates by Western blotting required that they first be concentrated by immunoprecipitation. In our imaging studies, we reproducibly observed stronger fluorescent signals from Vps8a, c, and d, compared to Vps8b, e, and f, particularly in live cell imaging. The weak fluorescence for Vps8e correlates with its apparent partial proteolytic cleavage (Fig. S2A, sixth lane), although proteolytic cleavage may occur during immunoprecipitation, rather than within live cells.

**Figure 3.**
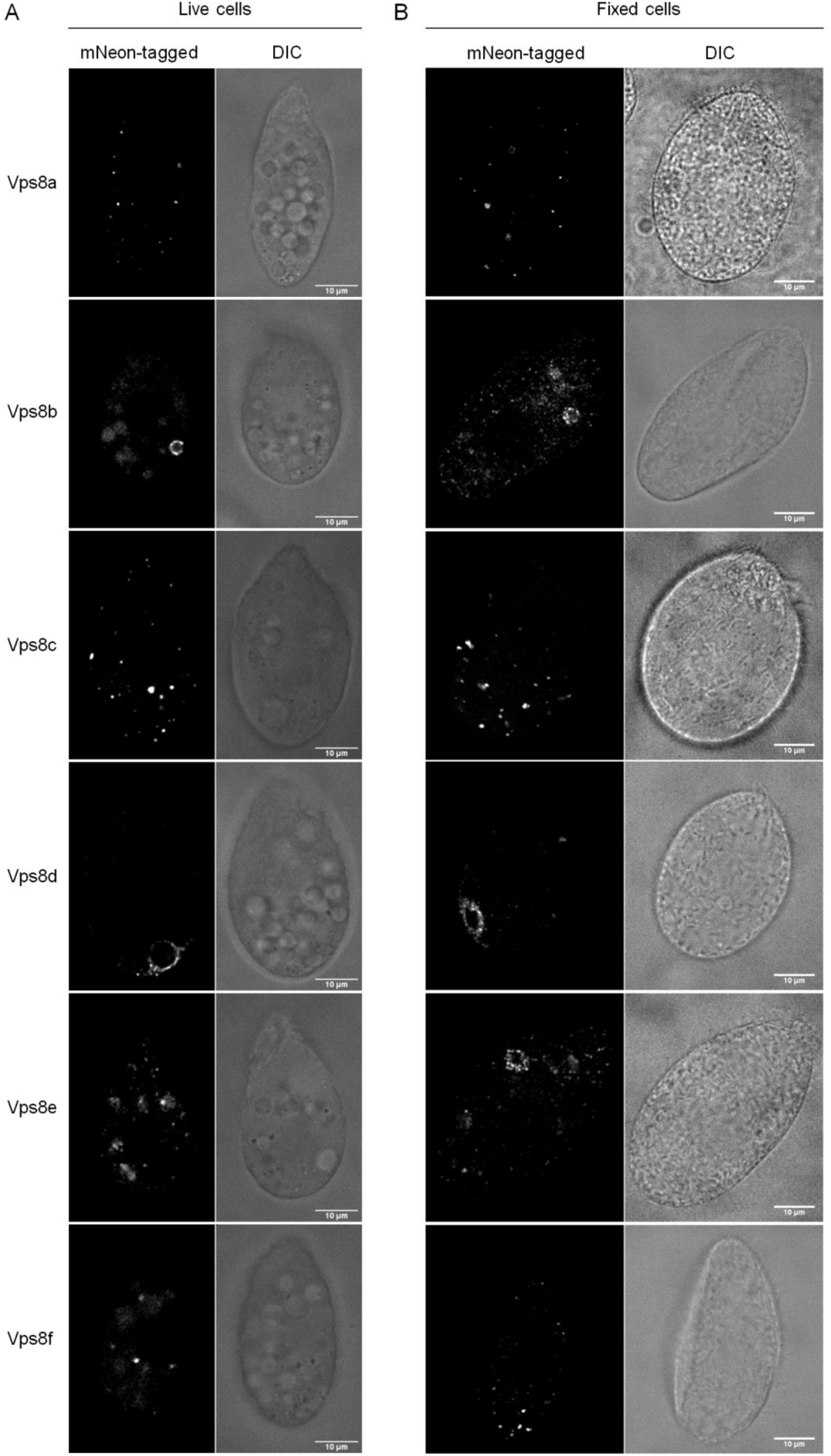
Each Vps8 paralog localizes to distinct cellular compartments. A) Live cell imaging of mNeon-tagged Vps8 paralogs revealed their primary distributions. Vps8a, as previously reported, localizes to small cytoplasmic vesicles. Vps8b localizes to vesicles at the periphery of large phagosomes/phagolysosomes. Vps8c localizes to cytoplasmic vesicles that are larger than those associated with Vps8a and to tubulovesicular compartments, and is concentrated in the posterior half of cells. Vps8d localizes to the contractile vacuole. Vps8e localizes to small vesicles of a uniform size evenly distributed within the cell. Vps8f localizes to few cytoplasmic puncta. Shown for Vps8a, b, c, e and f are single frames from time-lapse videos, with paired differential interference contrast (DIC) images, of cells endogenously expressing the mNeon-tagged Vps8 paralog. For Vps8d, a single confocal image was selected from a z-stack. All cells were incubated in S medium for 2h prior to imaging. B) Confocal sections of fixed cells expressing mNeon-tagged Vps8 paralogs, with paired DIC images. Cells were maintained in SPP prior to fixation. The distribution of fluorescent puncta seen in live cells was generally similar to that in fixed cells, but with some differences. In fixed cells, Vps8b-mNeon was associated with phagolysosomes but also with free cytoplasmic vesicles. For Vps8e, fixed cell sections revealed a population of vesicles at the cell anterior, close to the oral apparatus. Scale bars, 10μm.

While fully delineating the localization of these Vps8 paralogs requires additional compartmental markers for *Tetrahymena* to be developed, we draw some important conclusions. Vps8a-mNeon associates with small heterogenous puncta, many of which are highly mobile, and correspond to transport vesicles since they also contain the Sor4 receptor(Sparvoli et al., 2018). The most closely related paralog, Vps8c, also localizes to cytoplasmic puncta, but these are larger and include irregular structures that appear, in some instances, to possess tubular extensions (Fig. 3A, B, third panels). At least one such tubulovesicular structure was seen in every cell, and they are more frequently located toward the posterior. This posterior bias was more obvious in fixed cells, probably because compression of cells under the coverslip reveals more structures in any given focal plane (figure 3B third panel).

Vps8b-mNeon fluorescence appears in both live and fixed cells as a tightly-spaced array of puncta at the periphery of large circular structures, whose size and shape mark them as probable food vacuoles (Fig. 3A, 2nd panel) (Fig. 3B, 2nd panel). In addition, isolated small cytoplasmic puncta are visible in fixed cells, and, to a lesser extent, in favorable focal planes of live cells (Fig. S2B). Vps8d-mNeon fluorescence is strikingly concentrated at the contractile vacuole, a tubulovesicular organelle localized toward the cell posterior that functions in osmoregulation (Fig. 3A, B, forth panel). Vps8e-mNeon fluorescence appears in numerous small puncta throughout the cell cytoplasm (Fig. 3A, B, fifth panel). However, in some fixed samples from cells in growth media, the puncta were concentrated around a single circular structure near the cell anterior, close to the oral apparatus where food vacuoles are formed (Fig 3B, fifth panel). Finally, Vps8f-mNeon puncta, though almost undetectable in live cells (Fig. 3A, sixth panel), are clearly visible in fixed cells (Fig. 3B, sixth panel). These puncta are highly heterogeneous, and show no obvious pattern. For live imaging of Vps8f and all Vps8 paralogs with weak signals, the long exposure times result in auto-fluorescent food vacuoles that are also seen in wildtype cells.

### Vps8b and 8e localize to food vacuoles at two different stages

*Tetrahymena* are avid bacterivores and rapidly concentrate bacteria *via* ciliary beating at the anterior-positioned oral apparatus. From the base of the oral apparatus bacteria are taken up into newly-formed food vacuoles and subsequently digested as food vacuoles mature(Nilsson, 1979). Based on their fluorescence patterns, Vps8b-mNeon and Vps8e-mNeon appear to localize to food vacuoles. To confirm this, we labeled food vacuoles by incubating *Tetrahymena* with dsRed-expressing *E. coli*. The circular structures associated with Vps8b and Vps8e both contain dsRed fluorescence (Fig. 4, upper and lower panel, respectively). In the case of Vps8e structures, little or no digestion had yet occurred since the bacteria appeared intact. Both this observation and the anterior position of these structures, suggest that Vps8e associates with newly-formed food vacuoles. In the Vps8b structures, in contrast, no intact bacteria were present, indicating that these are more mature food vacuoles.

**Figure 4.**
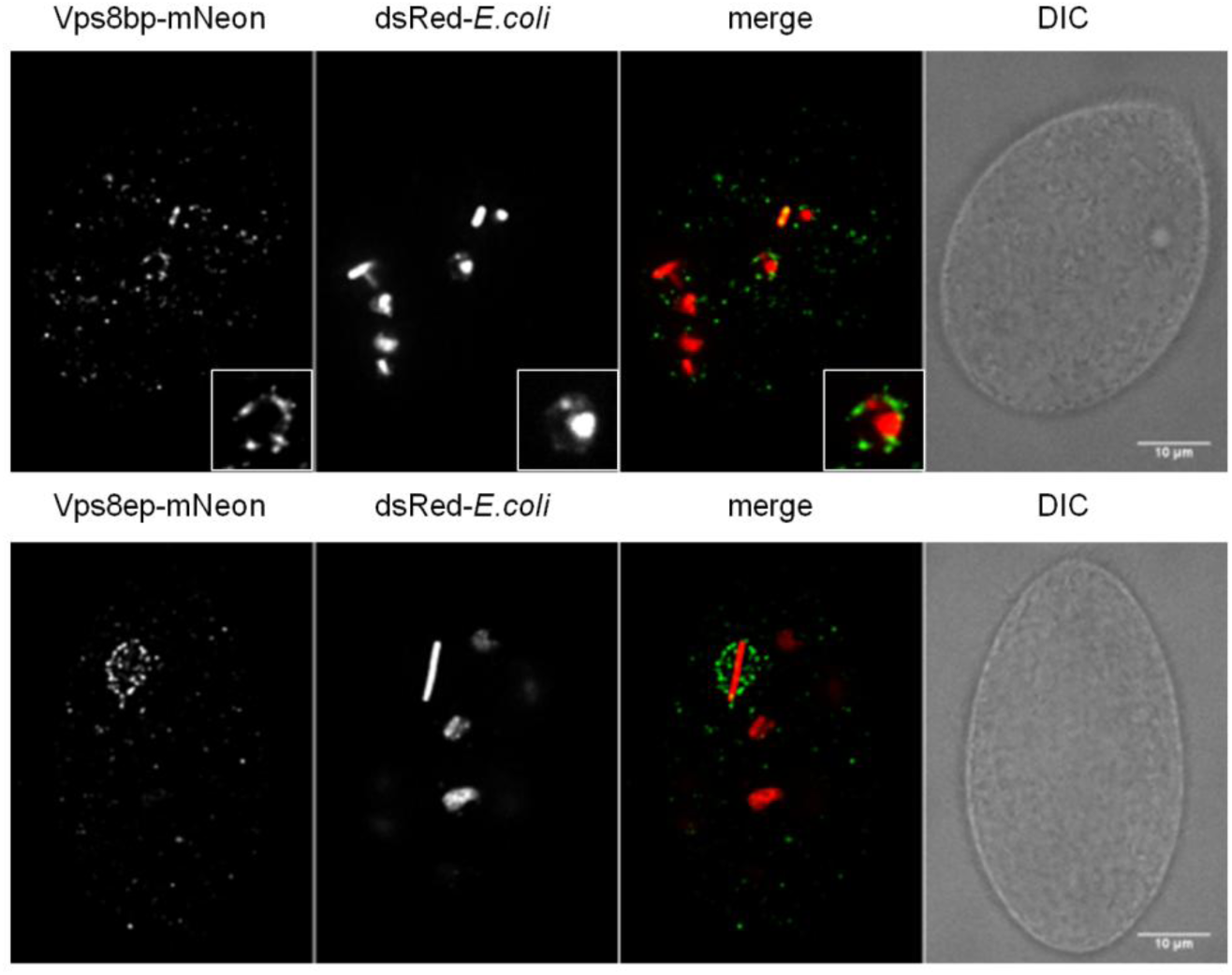
Localization of Vps8b and Vps8e to bacteria-containing phagosomes. Cells expressing Vps8b-mNeon (upper panel) and Vps8e-mNeon (lower panel), were fed with *E.coli* expressing dsRed for 30 min and then fixed. For Vps8b cells, the inset shows a structure similar to the Vps8b-labeled structures in figure 3, and which also contains dsRed bacteria. For Vps8e cells, the labeled anterior vesicles can be seen to surround an intact bacterium. Shown are confocal sections, with paired DIC images to the right. Scale bars, 10 μm.

### The evolutionarily related Vps8a and Vps8c, and by inference 8A-CC and 8C-CC, show very limited overlap

HOPS and CORVET in other lineages share four subunits, but they differ in their two Rab-binding subunits and therefore are targeted to different compartments(Solinger and Spang, 2013). Hybrid CORVET/HOPS complexes that differ from either HOPS or CORVET at only a single Rab-binding subunit have been detected in *S. cerevisiae*, but these complexes are of unknown significance(Peplowska et al., 2007). In considering the cohort of CORVET complexes in *Tetrahymena*, one striking inference is that the evolutionary replacement of the Vps8 subunit was sufficient to provide novel function, even when all other subunits remained identical. That is, although 8A-CC and 8C-CC are identical except Vps8, they localize to two seemingly different structures. The same is true for the 8E-CC and 8F-CC pair.

To understand this further we focused on the most recently diverged paralogs, Vps8a and 8c, and by inference 8A-CC and 8C-CC. Vps8a appeared to localize primarily to different structures than Vps8c, but we could not rule out the possibility for significant overlap in localization. To examine this possibility more rigorously, we generated cell lines in which complementary pairs of fluorescently-tagged CORVET subunits were simultaneously expressed. As a positive control, we created cells simultaneously expressing Vps8c-mNeon and Vps8c-mCherry. In such cells, there was the expected extensive overlap between the red and green signals (Fig. 5A, B). In our negative controls, we found the expected limited overlap when we co-expressed Vps8e-mCherry with either Vps8a-mNeon or Vps8c-mNeon (Fig.5C, upper and middle panels, respectively, and 5D). Significantly, cells simultaneously expressing Vps8a-mNeon and Vps8c-mCherry also showed very limited overlap (Fig. 5C, lower panel, and 5D). The expression of full-length fusions in each cell line was confirmed by SDS-PAGE and western blotting (Fig. S3A). This result indicates that the steady state localization of these CORVET complexes is primarily determined by Vps8, rather than any other subunit.

**Figure 5.**
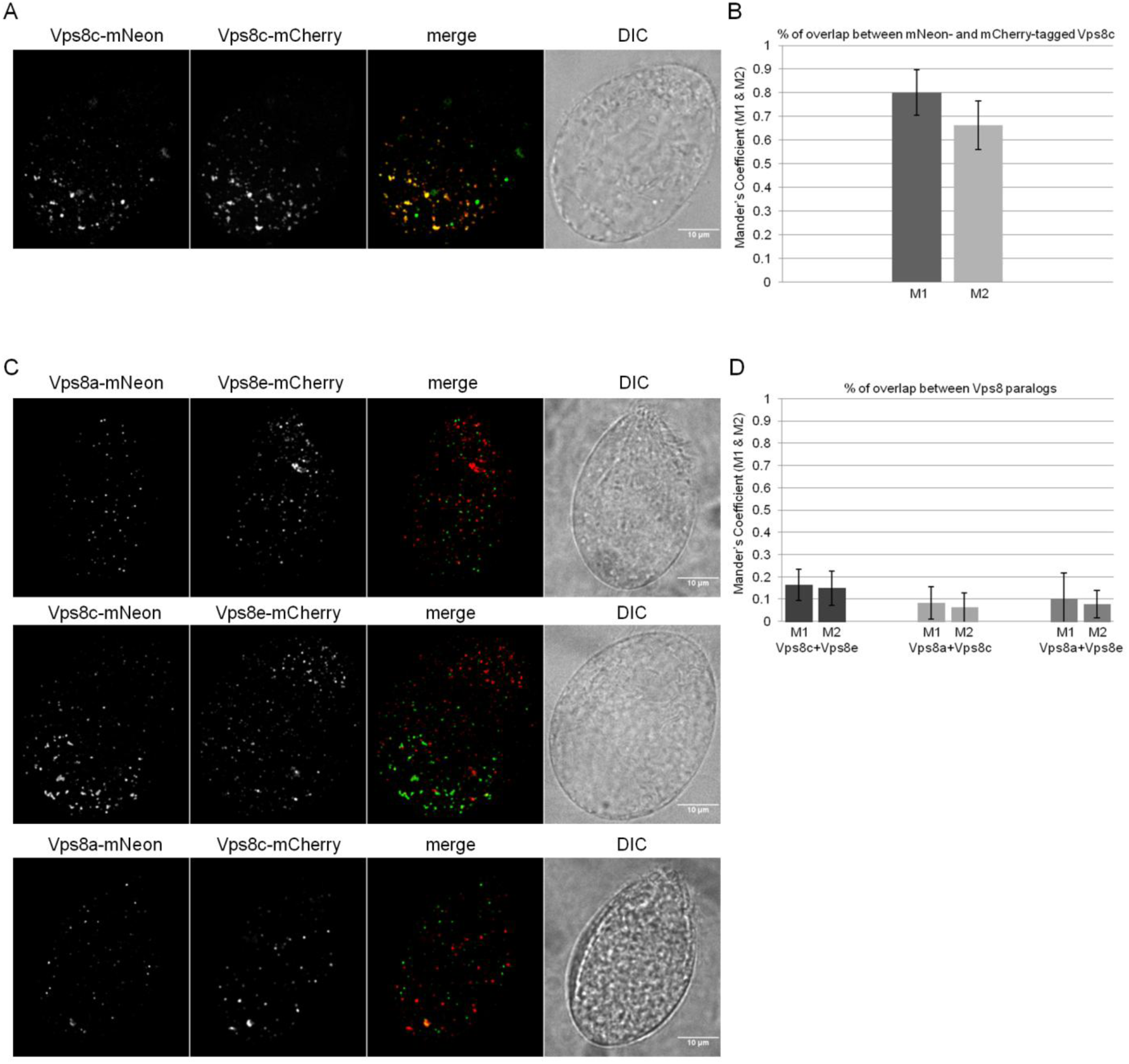
8A-CC and 8C-CC localize to non-overlapping compartments. A) Cells were transformed to simultaneously express Vps8c-mNeon and Vps8c-mCherry at the *VPS8C* and *VPS8A* loci, respectively. Shown is a maximum-intensity projection of a confocal z-stack of a fixed cell. Paired DIC images are confocal cross sections. B) Mean values of the co-localization coefficients M1 and M2 for Vps8c-mNeon and Vps8c-mCherry were calculated with 261 non-overlapping images using Fiji-JACoP plugin. Error bars represent Standard Deviations. The two fusion proteins largely overlap. C) Cells were transformed to simultaneously express the following combinations of tagged proteins: Vps8e-mCherry with Vps8a-mNeon (top); Vps8e-mCherry with Vps8c-mNeon (middle); Vps8a-mNeon with Vps8c-mCherry (bottom). All tagged proteins were integrated at their respective endogenous locus. Shown are maximum intensity projections of confocal z-stacks of fixed cells. D) Mean values of the co-localization coefficients M1 and M2 for Vps8a/Vps8e, Vps8c/Vps8e, and Vps8a/Vps8c pairs were calculated with 246/337/399 non-overlapping images/sample as in (B), respectively. Error bars represent Standard Deviations. The paralogs localize to largely non-overlapping cellular structures. Scale bars, 10 μm.

Vps8a-mNeon undergoes limited proteolytic cleavage in cells, as detected by Western blotting (Fig. S2A, second lane). We were concerned that the distribution of intact Vps8a-mNeon might not reflect the full distribution of Vps8a in cells. We took advantage of our previous finding that Vps8a tagged with GFP is functional and does not undergo proteolytic cleavage(Sparvoli et al., 2018) (Fig. S3B). We therefore compared the number of fluorescent puncta in Vps8a-GFP vs Vps8a-mNeon cells (Fig. S3C). Since there was no significant difference (Fig. S3D), we conclude that Vps8a-mNeon provides a suitable reporter for the localization of the endogenous protein, and that Vps8a and Vps8c are chiefly localized to non-overlapping structures.

To obtain additional evidence that these Vps8 proteins belong to full CORVET complexes, we tested the inference that the shared subunits Vps11 and Vps3a are present in diverse compartments labeled by the set of Vps8 paralogs. In cells that co-express Vps3a-GFP with Vps11-mCherry, at their endogenous loci, the proteins are highly co-localized (Fig. 6A, C). Additionally, as expected Vps3a co-localizes extensively with Vps8c (Fig. 6B, C). Most importantly, cells individually expressing either GFP-tagged Vps3a or Vps11 labeled a diverse set of structures, including cytoplasmic vesicles of different sizes, phagolysosome-related structures similar to those labeled by Vps8b (only for Vps11-GFP) and Vps8e, and the contractile vacuole (Fig. 6D, E).

**Figure 6.**
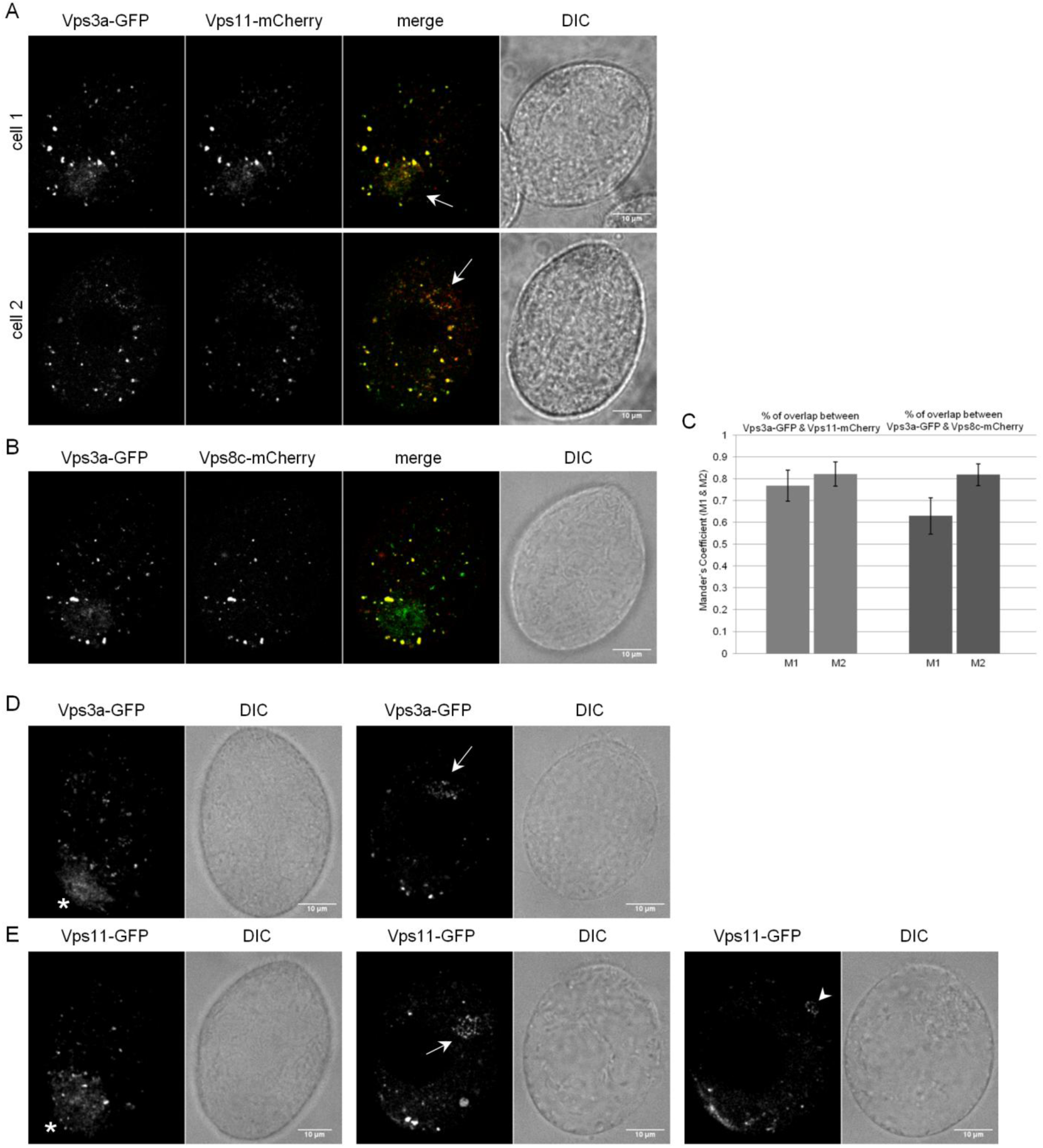
Vps3a and Vps11 localize to a wider range of compartments than individual Vps8 subunits. A) Confocal sections of cells co-expressing Vps3a-GFP and Vps11-mCherry. The two subunits largely colocalize at heterogeneous cytoplasmic vesicles, including vesicles close to the oral apparatus (white arrow in cell 2) resembling those associated with Vps8e, and at the contractile vacuole (white arrow in cell 1). B) Confocal section of cells co-expressing Vps3a-GFP and Vps8c-mCherry. The two subunits significantly overlap on heterogeneous cytoplasmic vesicles. In addition, Vps3a-GFP alone is visible at some structures, including the contractile vacuole. C) Percentages of overlap (Mander’s coefficients M1 and M2) for Vps3a/Vps11 and Vps3a/Vps8c were derived from 325 and 321 non-overlapping images using Fiji-JACoP plugin, respectively. Vps3a extensively overlaps with both Vps11 (left columns) and Vps8c (right columns). Error bars represent Standard Deviations. D) and E) Confocal sections of cells expressing Vps3a-GFP or Vps11-GFP. The overall distributions of Vps3a and Vps11 largely recalls that of the combination of individual Vps8 paralogs, and include heterogeneous cytoplasmic vesicles, the contractile vacuole (asterisk), and rings of vesicles near the oral apparatus like those labeled by Vps8e (arrow). Vps11 but not Vps3a is also found at rings of vesicles around food vacuoles (arrowhead), like those labeled by Vps8b. The images showing contractile vacuoles are all surface sections, while the others are cell mid sections. DIC images are shown to the right of each panel. Scale bars, 10 μm.

### Vps8a and Vps8c localize to non-equivalent Rab7-labeled compartments

In other organisms CORVET acts at Rab5-positive compartments, while HOPS functions at Rab7-positive compartments. Rab5 was lost in the *Tetrahymena* lineage, but *T. thermophila* expresses the related Rab22a (Bright et al., 2010). We previously reported the surprising finding that Vps8a shows negligible co-localization with Rab22a but ∼50% co-localization with Rab7(Sparvoli et al., 2018). To determine if this was also the case for Vps8c, we expressed N-terminal mCherry-tagged Rab7 (Fig. 7A) and Rab22a (Fig. 7B) in Vps8c-mNeon expressing cells, and measured the extent of overlap. Vps8c co-localized more strongly with both Rabs than Vps8a (Fig. 7C, D), and in particular ∼90% co-localization with Rab7 (Fig. 7C). This co-localization could also be seen at dynamic tubulo-vesicular structures (Fig. S4A, B). Since Vps8a and Vps8c both significantly co-localize with Rab7, but very little with one another, there must be additional localization determinants present.

**Figure 7.**
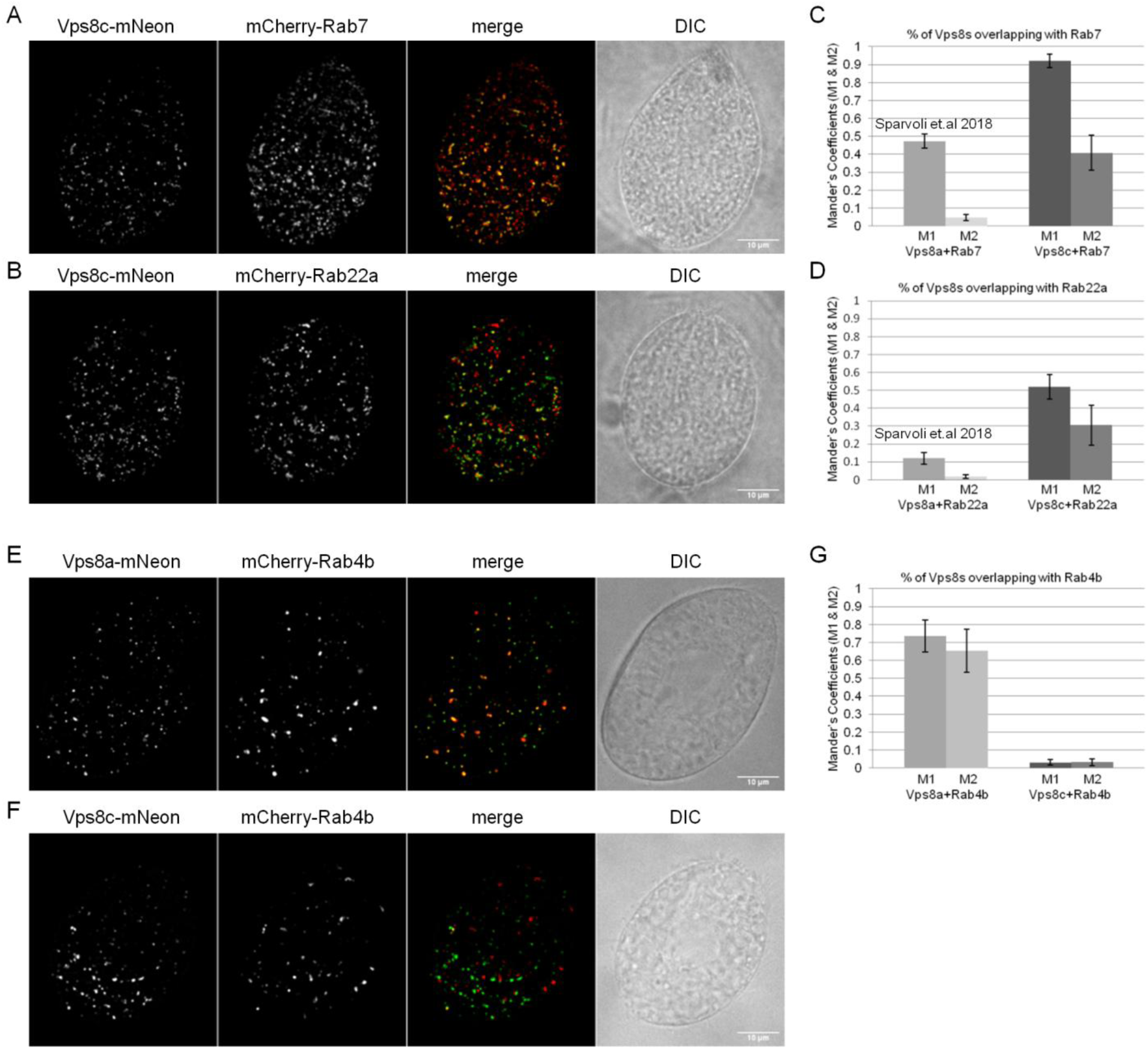
8A-CC and 8C-CC associate with distinct endosomes. A) and B) Cells were transformed to co-express Vps8c-mNeon at the endogenous locus together with either mCherry-Rab7 (upper panel) or the Rab5 homolog mCherry-Rab22a (lower panel). Vps8c-mNeon was expressed at the *VPS8C* locus, while Rab transgenes were expressed from the *MTT1* promoter by induction with 1 μg/ml CdCl_2_ for 2 h in SPP. Shown (including E, F) are maximum intensity projections of z-stacks of fixed cells. The DIC images are confocal cross sections for clarity. C) and D) Percentages of overlap (Mander’s coefficients M1 and M2) for Vps8c/Rab7 and Vps8c/Rab22a were calculated using Fiji-JACoP plugin. The Mean M1 and M2 values and Standard Deviations (error bars) for Vps8c were derived from 268 and 232 non-overlapping images for Rab22a and Rab7 samples, respectively. E) and F) Cells were transformed to endogenously co-express Vps8a-mNeon (upper panel) or Vps8c-mNeon (lower panel) with mCherry-Rab4b. G) Percentages of overlap (Mander’s coefficients M1 and M2) for Vps8a/Rab4b, and Vps8c/Rab4b, were calculated with 158/139 non-overlapping images/sample, respectively, using Fiji-JACoP plugin. In contrast with Vps8c-mNeon, Vps8a-mNeon shows extensive co-localization with Rab4b. Error bars represent Standard Deviations. Scale bars, 10 μm.

We sought other Rab proteins that could act as determinants in the Vps8a-dependent pathway of mucocyst formation. We previously used a genome-wide approach to identify genes upregulated in periods of stimulated mucocyst formation(Haddad et al., 2002)(unpublished). By searching these data for Rabs we identified a Rab4/Ypt31homolog (Rab4b; (Bright et al., 2010)), that is induced 11-fold under conditions of induced mucocyst formation. Moreover, the transcriptional profile of the *RAB4B* gene, under a variety of cell culture conditions, is strikingly similar to a variety of established mucocyst-associated genes, suggesting these genes are co-regulated (Fig. S4C). To determine if Rab4b might contribute towards Vps8a localization, we endogenously tagged Rab4b with mCherry at its N-terminus, and co-expressed in cells with Vps8a-mNeon or Vps8c-mNeon (Fig. 7E and F, respectively). Strikingly, Vps8a, but not Vps8c, showed strong co-localization with Rab4b (Fig. 7G).

Taken together, our results suggest that all six Vps8 paralogs, and by inference their parent CORVET complexes, are individually specialized for distinct trafficking pathways in *Tetrahymena*. Specialization extends to even the most recently diverged paralogs: 8A-CC and 8C-CC, which can be considered sibling complexes, are largely non-overlapping in their distribution. 8C-CC shows significant co-localization with early endosomal Rab22a, and almost complete co-localization with late endosomal Rab7. 8A-CC, which co-localizes partially with Rab7, has very strong co-localization with Rab4b, which is a recycling endosome marker.

## Discussion

The loss of the HOPS tethering complex was accompanied in a sub-family of ciliates by expansions in the number of CORVET complex subunits. In *T. thermophila* we report that there are six biochemically distinct CORVET complexes, which we call 8A-CC, 8B-CC, etc., each possessing a different Vps8 paralog. While detailed functional analysis has yet to be achieved our localization data strongly suggest that the complexes diversified to associate with six distinct compartments. That is, five of the six endogenously-tagged Vps8 paralogs localize to structures that are recognizably different from one another in live cells. Consistent with the idea of functional specialization, there is differential expression of genes expressing subunits that are specific to individual complexes, as seen in whole genome transcriptional profiling over a range of culture conditions.

The CORVET complexes are expressed at very low levels, but we could nonetheless isolate all six complexes by taking advantage of cryomilling that has been used in other organisms. Determining the subunit composition of all six complexes allowed us to make some observations about the pattern of diversification within CORVET genes. In two pairs of complexes the only difference is the Vps8 subunit and for 8A-CC and 8C-CC there is minimal co-localization. CORVET targeting depends upon Rab protein interactions *via* the two Rab-binding Vps8 and Vps3 subunits (Epp and Ungermann, 2013). In *Tetrahymena,* at least, the Rab-binding by these subunits is not functionally equivalent; specifically since 8A-CC and 8C-CC share Vps3a, they would be expected to co-localize if Vps3 were a primary targeting determinant. Since this is not observed, the Vps8 subunit is likely primarily responsible for the differential steady-state location of most CORVET complexes. This idea is broadly consistent with experiments in yeast HOPS, showing that the Rab-binding subunits Vsp39 and Vps41 have different binding properties(Lurick et al., 2017). Vsp39 and Vps41 are positioned at opposite ends of the extended barbell-like cryoEM structure of yeast HOPS, consistent with independent binding and with the idea that relative Rab affinities could determine steady state localization(Brocker et al., 2012). A current assumption is that CORVET and HOPS have similar overall structures, given the molecular similarities, but this remains to be demonstrated. The barbell structure was determined from cross-linked complexes using cryoEM, while analysis of non-cross-linked complexes suggests considerably flexibility, with the barbell just one of several relevant structures(Chou et al., 2016; Kuhlee et al., 2015). Therefore, a second and non-mutually exclusive potential explanation of our results is that alternative conformations for CORVET bias Rab binding in favor of Vps8.

In either model, the non-colocalization of different CORVETs bearing the identical Vps3a subunit could be explained if binding of Vps3 to its cognate Rab is rapidly followed by membrane fusion, i.e., bivalent tethering is a short-lived state. Alternatively, non-colocalization could be explained if Vps3a recognizes a Rab present on a variety of membranes, e.g., sub-compartments of an organelle. In this scenario, additional contacts might refine the targeting of each CORVET complex to an individual sub-compartment. We observed that although both 8C-CC and 8A-CC individually colocalize with Rab7, discussed further below, they show minimal co-localization with one another. This suggests that multiple distinct populations of Rab7-positive endosomes are present, similar to distinct sub-populations of early endosomes in mammals(Perini et al., 2014). Consistent with this idea, *Tetrahymena* Rab7 labels highly mobile cytoplasmic vesicles but also the contractile vacuole and phagosome-associated vesicles (unpublished data), the last consistent with biochemical data(Jacobs et al., 2006). It is thus possible that Rab7 is present at all CORVET-positive structures.

In yeast and animals, CORVET or HOPS subunits also engage in non-Rab-based interactions with target membranes(Fratti et al., 2004; Ho and Stroupe, 2016; Stroupe et al., 2006). Similarly, combinatorial interactions could explain how different CORVET complexes in *Tetrahymena* are differentially recruited to Rab7-positive membranes. In this regard, it is notable that several CORVET subunit paralogs in *Tetrahymena* are larger than their homologs in Opisthokonts, potentially offering novel structures for interactions. Vps8a and 8e are 22% and 58% larger, respectively, than their *S. cerevisiae* ortholog. Size variation for Vps8 is also notable in plants where *Arabidopsis* Vps8 is 46% larger than in yeast. By contrast Vps11, a subunit with an organizing role in assembly of the HOPS/CORVET core(Ostrowicz et al., 2010; Plemel et al., 2011), has remained nearly invariant in size suggesting significant selective pressure for conservation.

An implication of CORVET composition in *Tetrahymena* is that most tethers are not acting as homotypic tethers, unlike CORVET in other organisms that have been analyzed. That is, in the five complexes containing Vps3a, the five different Vps8 paralogs are unlikely to all share the Vps3a Rab-binding specificity. As heterotypic tethers, the *Tetrahymena* complexes may shed light on hybrid CORVET/HOPS complexes that have been reported in yeast, which have the potential to bind both Rab5 and Rab7(Peplowska et al., 2007). These hybrids have been hypothesized to represent intermediates in CORVET-to-HOPS switching during endosome maturation, in a mechanism involving step-wise substitution of complex-specific subunits on the shared core, and discussed further below. However, we note that such hybrid CORVET/HOPS tethers have to date only been detected in yeast, and under conditions in which subunit overexpression could potentially result in non-physiological complexes.

The pattern of subunit variation between complexes suggests that the core Vps18 subunit may determine which Vps8 paralog is included. Among Vps8 paralogs, Vps8a and 8c are relatively closely related, as are Vps8e and f, while Vps8b and 8d are more divergent from the others(Sparvoli et al., 2018). Based on data here, the cores containing Vps18d form complexes that also contain Vps8a or Vps8c, while cores containing Vps18c instead assemble with Vps8e or 8f subunits. The single cores containing Vps18a or Vps18b assemble with the highly unrelated Vps8d and Vps8b subunits, respectively. The idea that Vps18 paralogs determine the inclusion of specific Vps8 paralogs is consistent with genetic, biochemical and structural mapping of subunit interactions (Guo et al., 2013; Hunter et al., 2017; Plemel et al., 2011) (Brocker et al., 2012; Chou et al., 2016).

The function of 8A-CC was analyzed in previous work(Sparvoli et al., 2018), while results in this paper provide hints about functions of the other five CORVET complexes. At least two are associated with the pathway of phagocytosis and food vacuole formation. 8E-CC is targeted to vesicles associated with phagosomes, which are likely to be newly-forming based on their anterior position. It may be required for their tethering and fusion, since *Δvps8e* cells accumulate an excess of small endocytic vesicles(Sparvoli et al., 2018). 8B-CC is associated with vesicles found at the periphery of phagolysosomes at a later stage in the pathway. Interestingly, we observed a notable accumulation of endocytic vesicles around phagosomes in *Δvps8b* cells(Sparvoli et al., 2018), suggesting the existence of a class of endosomes that require 8B-CC for fusion but not for docking. The function of 8F-CC is not apparent from its localization. However, in preliminary experiments we saw strongly similar phagocytosis defects in *Δvps8e* and *Δvps8f* cells, suggesting that 8F-CC is also required for early steps in phagocytosis.

8D-CC is associated with the contractile vacuole, a Rab11a-positive (and therefore endolysosomal) organelle which is nonetheless not known to be involved in trafficking of endocytic or secreted proteins(Allen, 2000; Bright et al., 2010; Plattner, 2015). Contractile vacuole activity in *Tetrahymena* is based on repeated cycles of membrane fusion and fission between tubules and a central bladder, and 8D-CC may contribute to that process. Interestingly, contractile vacuole CORVET possesses the largest number of exclusive subunits (4/6) among the *Tetrahymena* complexes, including a unique paralog of the SNARE-binding Vps33 subunit. The second relatively unique complex, with three exclusive subunits, is 8B-CC. Interestingly, all six subunits defining 8D-CC were strongly detected by mass spectrometry, but the Vps8d subunit was significantly more abundant than the other five based on analysis by silver staining. One possibility is that Vps8d partially exists as a monomer, for which we have some preliminary evidence. HOPS and CORVET sub-complexes have been identified in other organisms, as outlined earlier, and may be highly lineage-specific. One important source of ambiguity in characterizing subcomplexes using biochemical approaches is that a substantial fraction of CORVET remains membrane-attached under non-denaturing conditions.

Our understanding of Rab-binding specificities of CORVET in *Tetrahymena* is limited to 8A-CC and 8C-CC. In yeast and animals, CORVET binds early endosomal Rab5/Vps21 via Vps3 and Vps8 subunits (Balderhaar et al., 2013; Epp and Ungermann, 2013; Markgraf et al., 2009; Peplowska et al., 2007), while HOPS subunits Vps41 and Vps39 bind late endosomal Rab7/Ypt7 (Plemel et al., 2011; Wurmser et al., 2000). These specificities are maintained in plants (Takemoto et al., 2018) but not *Tetrahymena*, since the CORVET subunit Vps8a associates with Rab4b, a marker for recycling endosomes, and with Rab7 rather than the Rab5-homolog Rab22a. Interestingly, Vps8 in Hela cells similarly associates with Rab4, as part of a sub-complex that provides a function distinct from holo-CORVET(Jonker et al., 2018). Although the overall organization of endosomal trafficking in *Tetrahymena* remains to be analyzed, Rab22a is likely to be a *bona fide* early endosomal marker(Bright et al., 2010). We found, relative to Vps8a, that Vps8c co-localizes even more strongly with Rab7, but in addition co-localizes with Rab22a and not Rab4b.

In yeast and animals, the mechanism of endosome maturation involves CORVET-dependent recruitment of a guanine nucleotide exchange factor (GEF) that activates Rab7, which then recruits HOPS(Nordmann et al., 2010). By this mechanism, Rab5-to-7 conversion during endosome maturation is linked with the engaging of successive tethers. This mechanism also exists in plants, although recent evidence suggests that the HOPS function in some plant pathways does not depend upon this kind of GTPase switch(Takemoto et al., 2018). In *Tetrahymena*, our results beg the question of whether there is maturation-linked switching between distinct CORVET complexes like 8C-CC and 8A-CC. In both Opisthokonts and Archaeplastids, the Rab-switching complex that links CORVET and HOPS is the Mon1-Ccz1 heterodimer, where Ccz1 possesses GEF activity(Cui et al., 2014; Kiontke et al., 2017; Nordmann et al., 2010). Interestingly, while *Tetrahymena* has an unambiguous Mon1 homolog, there is no convincing ortholog for Ccz1 (unpublished). Thus, together with the loss of HOPS in this lineage, Ccz1 may also have been lost.

The expansion and functional diversification of CORVET in ciliates provides a potent example of mechanisms underlying new trafficking pathways. Loss of HOPS in the lineage leading to *Tetrahymena* may indicate a simplification of pre-existing transport pathways. Such reductions in pathway complexity, relative to that present in the Last Eukaryotic Common Ancestor (LECA), are supported by a collapse in diversity of pan-eukaryote Rab and TBC-GAPs (Elias et al., 2012; Gabernet-Castello et al., 2013). We hypothesize that expansion of lineage-specific Rab proteins, which account for ∼40% of the repertoire (RabX in (Elias et al., 2012)), likely reflects subsequent pressures in ciliates that also drove diversification of CORVET-mediated endosomal pathways, but which could only be met by expansions of CORVET subunits. This is fully consistent with the organelle paralogy model proposed earlier whereby expansions in trafficking gene families facilitate the emergence of new pathways(Dacks and Field, 2007), and is also reminiscent of the evolution of multiple early endosomal pathways in many lineages, including yeasts and kinetoplastida, where Rab5 paralogs have arisen independently of each other. We suggest that a ‘backfilling’ mode, involving expansion following loss, is a potentially underappreciated aspect of trafficking diversity, especially as a number of key trafficking proteins are frequently, and independently, lost from multiple lineages.

## Materials and Methods

### Cell culture

*Tetrahymena thermophila* strains used in this work are in Table 2. Cells were grown overnight in SPP (2% proteose peptone, 0.1% yeast extract, 0.2% dextrose, 0.003% ferric-EDTA) supplemented with 250 ug/ml penicillin G, 250 ug/ml streptomycin sulfate, and 0.25 μg/ml amphotericin B fungizone, to medium density (1-3 x 10^5^ cells/ml). For biolistic transformation, growing cells were subsequently starved in 10 mM Tris buffer, pH 7.4, for 18-20 hours. Fed and starved cells were both kept at 30 °C with agitation at 99 rpm, unless otherwise indicated. For live microscopy, cells were transferred to S medium (0.2% yeast extract, 0.003% ferric-EDTA) for 2 hours prior to imaging. Culture densities were measured using a Z1 Coulter Counter (Beckman Coulter Inc.).

**Table 2.**
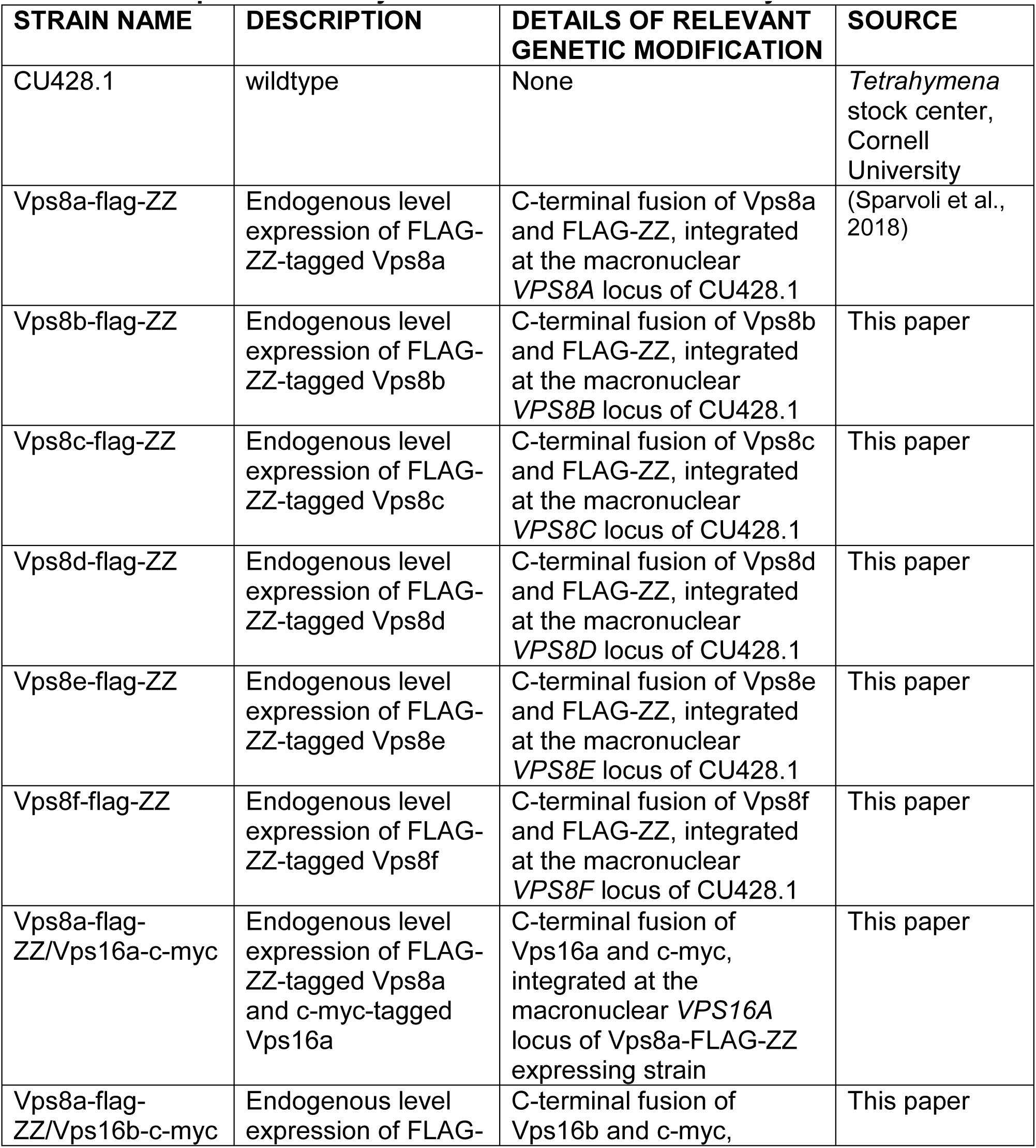
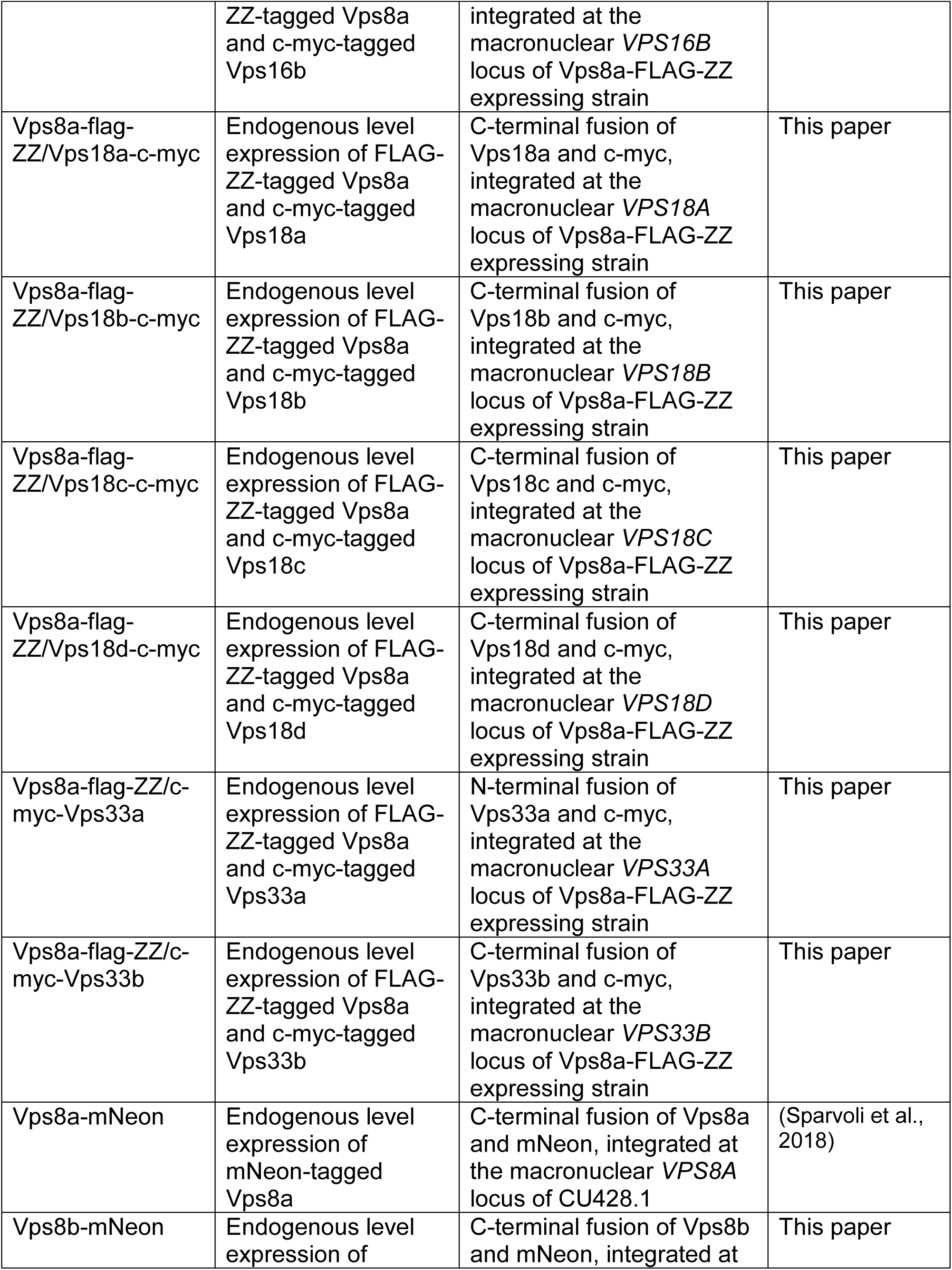
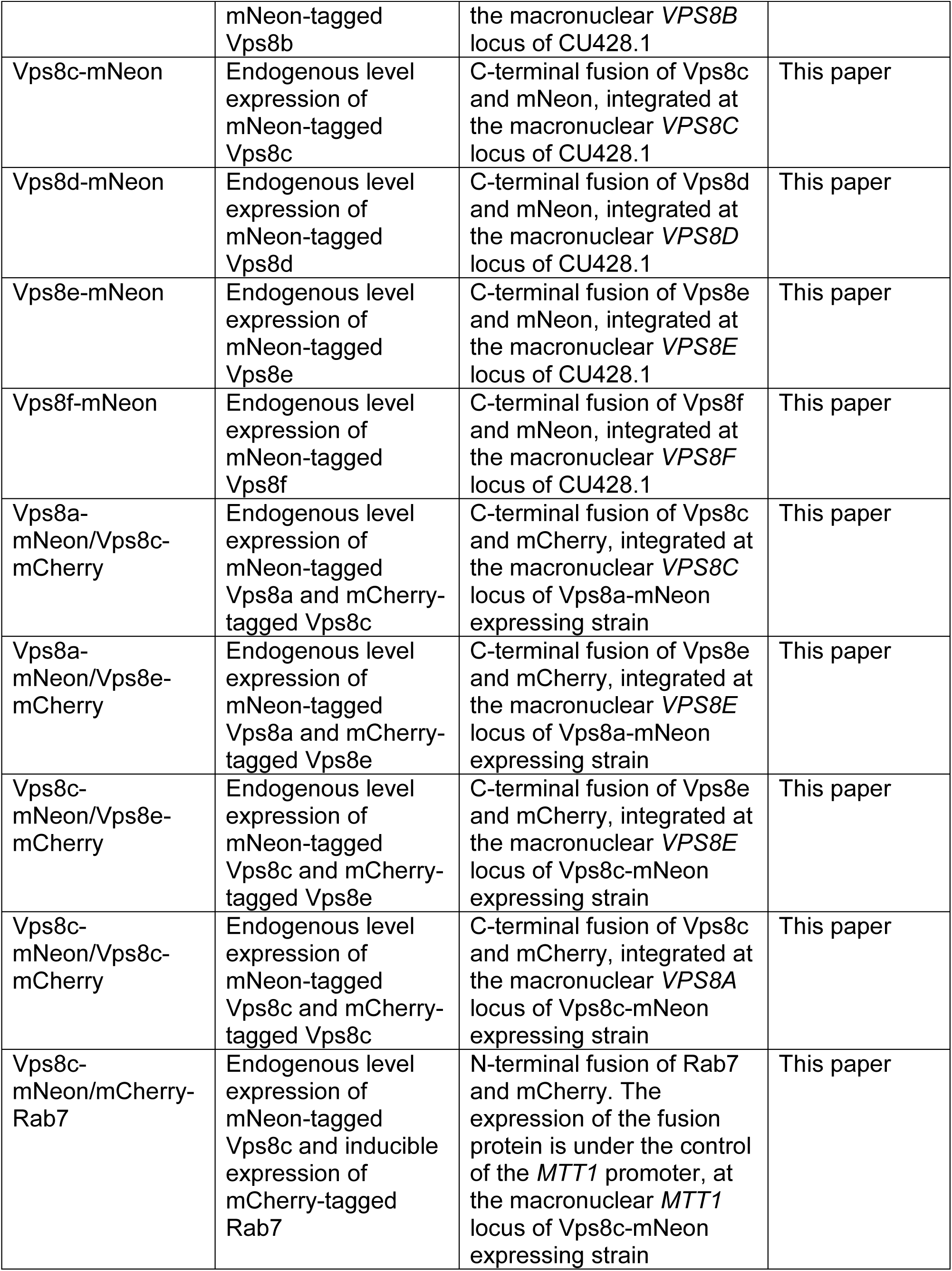
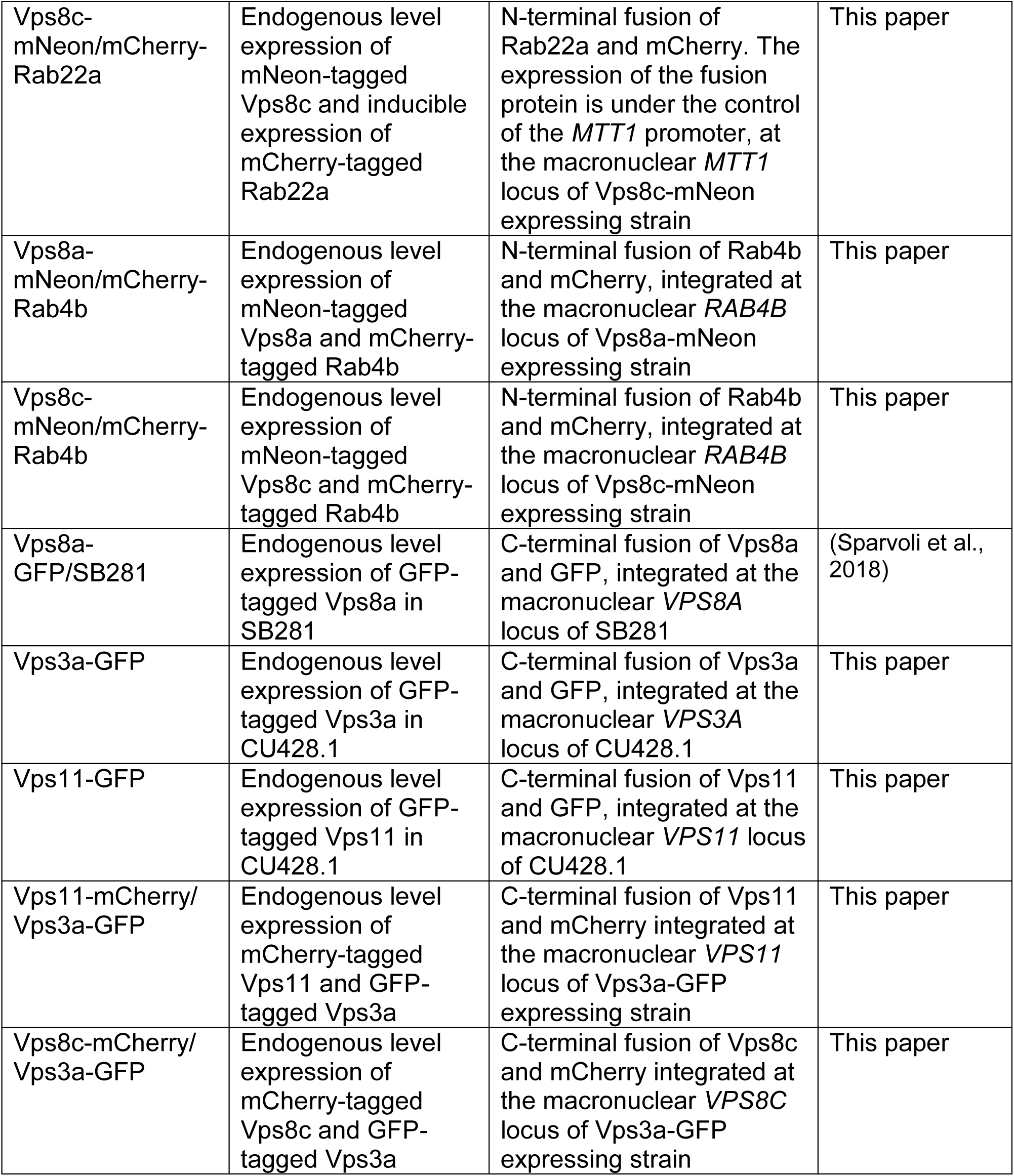
Description of Tetrahymena strains used in this study

### Endogenous tagging of the Vps8 paralogs with mNeon fluorescent tags

Two mNeonGreen fluorescent tags were integrated at the C-termini of the macronuclear ORFs of *VPS8B*, *VPS8C*, *VPS8D*, *VPS8E*, and *VPS8F* via homologous recombination using linearized pVPS8B-2mNeon-6myc-Neo4, pVPS8C-2mNeon-6myc-Neo4, pVPS8D-2mNeon-6myc-Neo4, pVPS8E-2mNeon-6myc-Neo4, pVPS8F-2mNeon-6myc-Neo4. The C terminal 745bp, 763bp, 763bp, 745bp, 468bp of *VPS8B*, *VPS8C*, *VPS8D*, *VPS8E*, *VPS8F* genomic locus (minus the stop codon), were amplified by PCR and cloned in digested p2mNeon-6myc-Neo4 vector (Sparvoli et al., 2018) at the SacI/MluI sites by Quick Ligation (New England, Biolabs Inc.), respectively. Subsequently, the 758bp, 744bp, 799bp-long 3’ UTRs of *VPS8B*, *VPS8C*, *VPS8D* were cloned in the VPS8-specific p2mNeon-6myc-Neo4 vector at the XhoI/ApaI, while the 808bp, 793bp 3’ UTRs of *VPS8E* and *VPS8F,* were cloned at the EcoRV/XhoI sites. The final constructs were digested with SacI and KpnI prior to biolistic transformation of CU428.1. The primers are listed in Table 3.

**Table 3.**
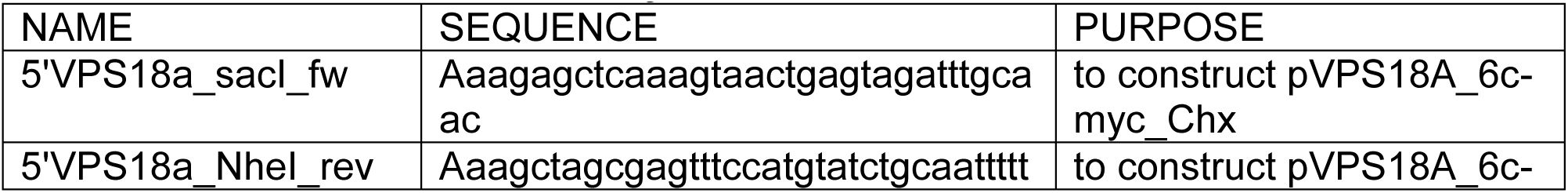
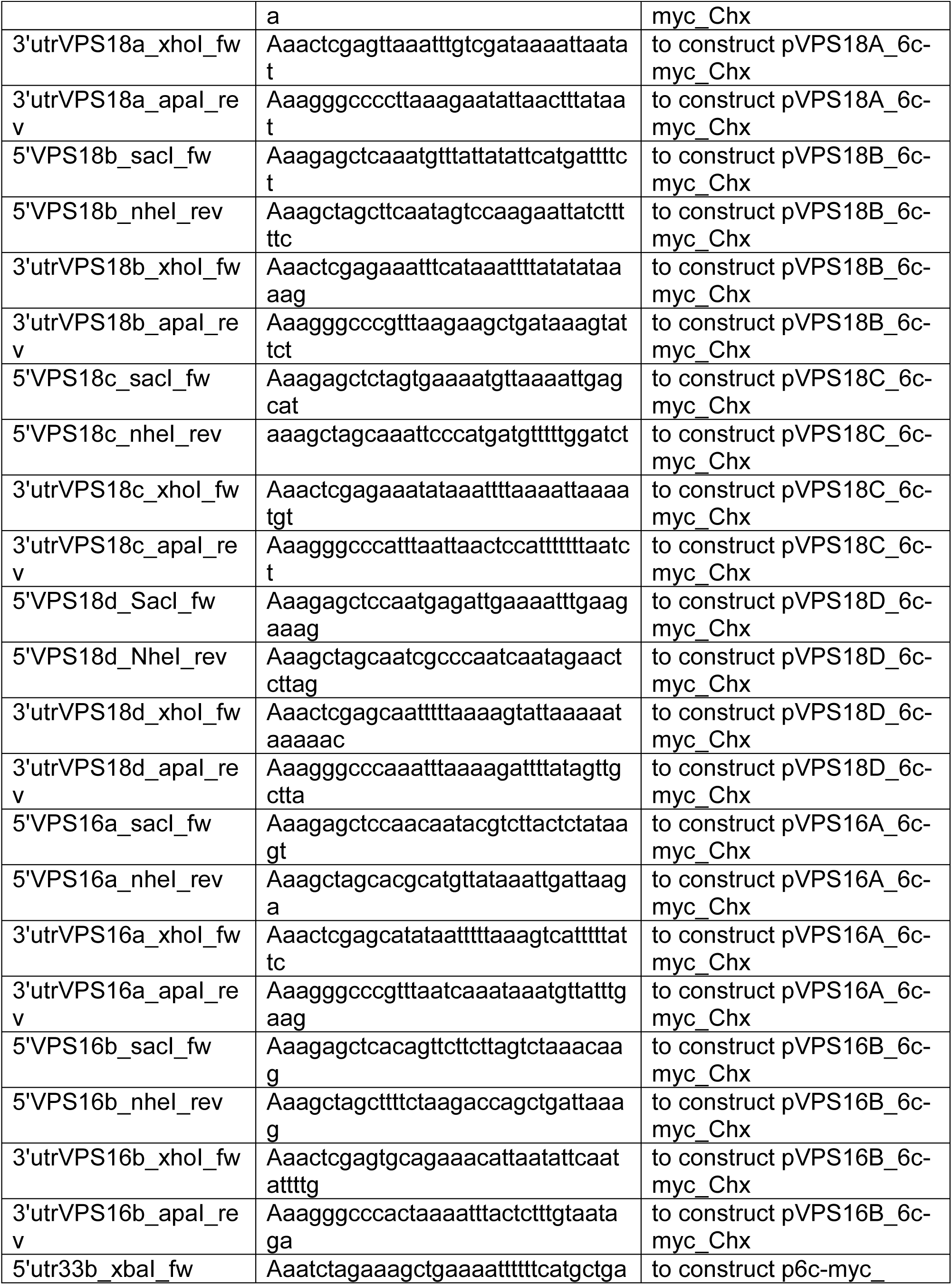
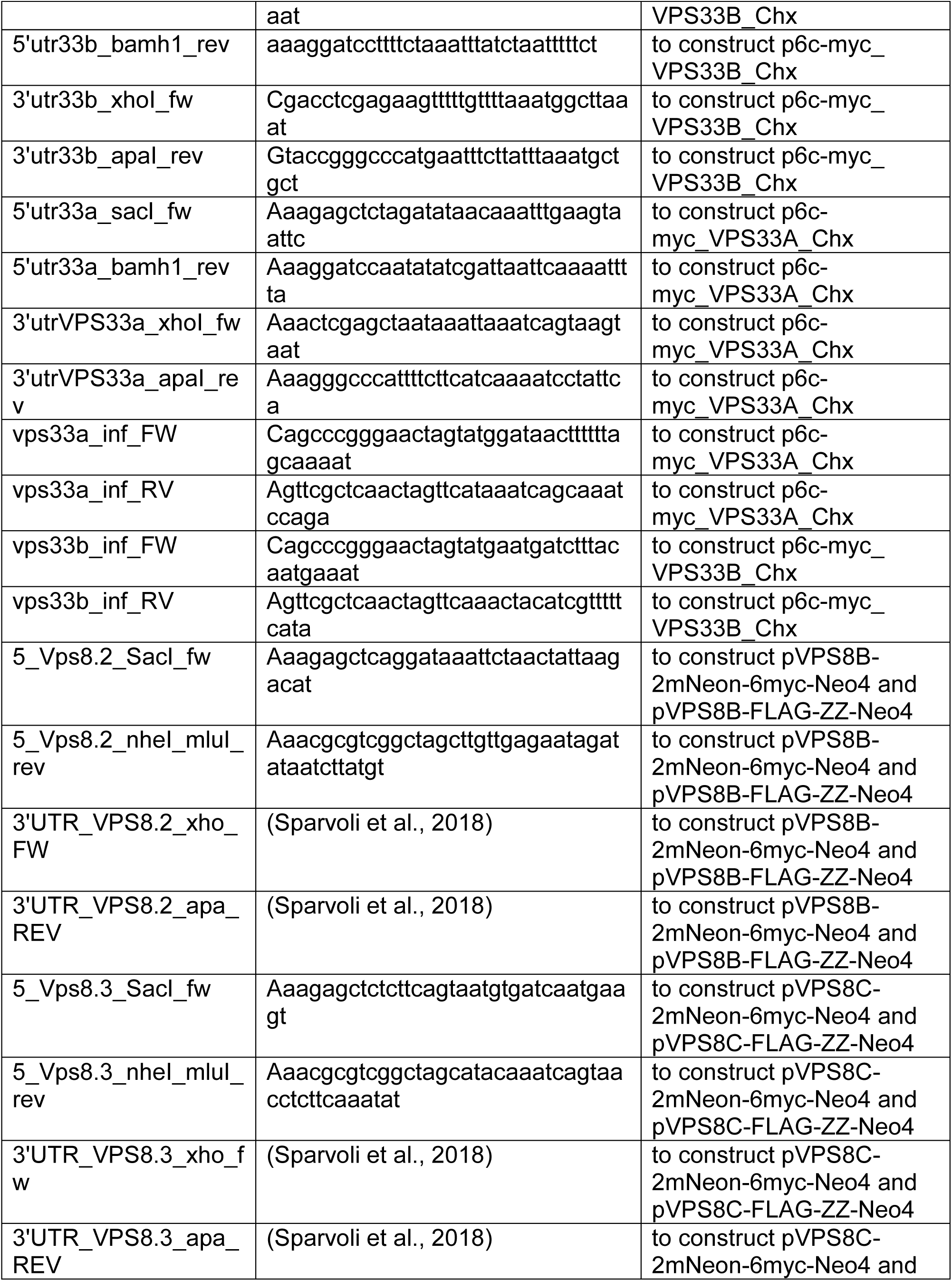
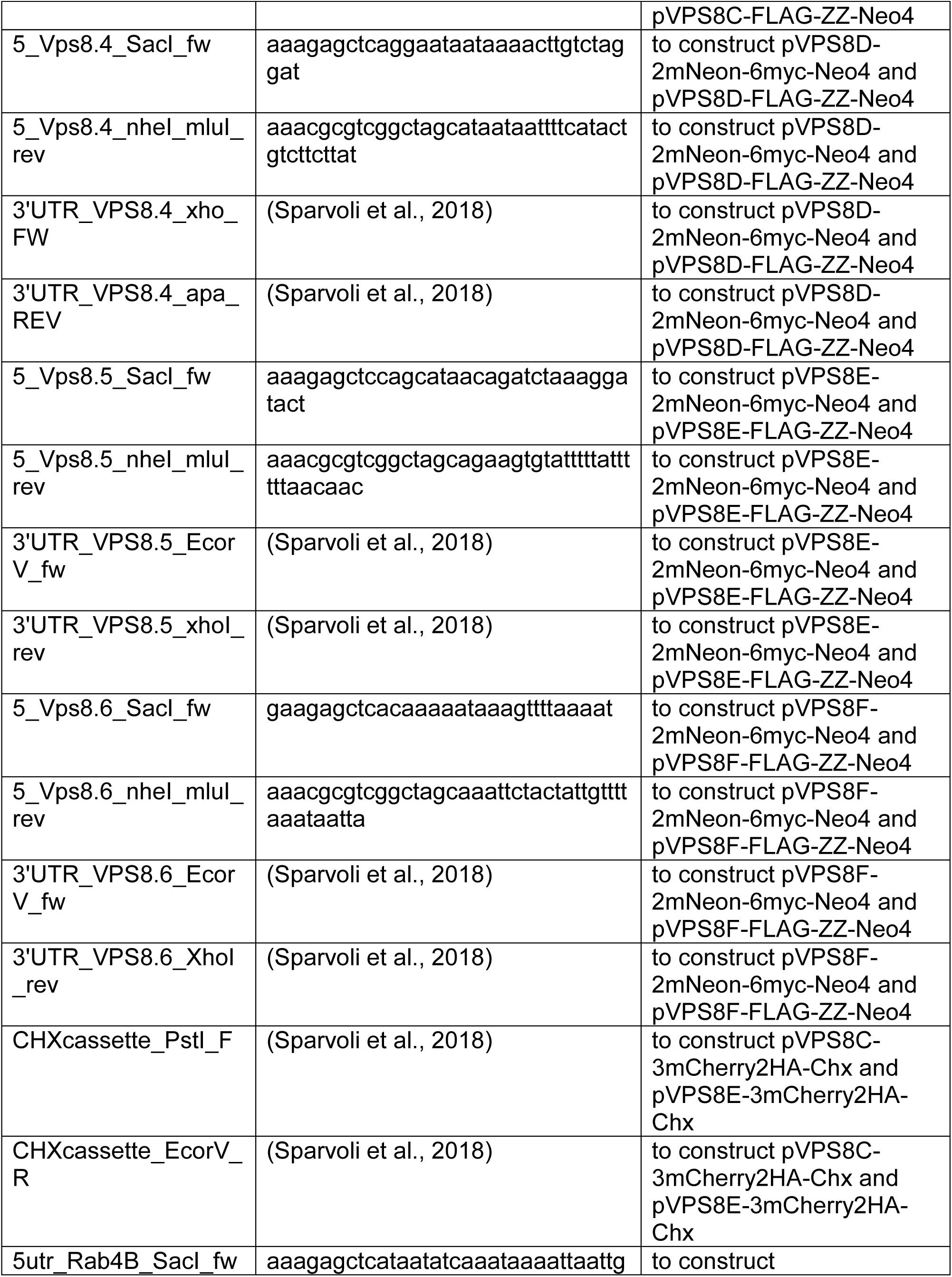
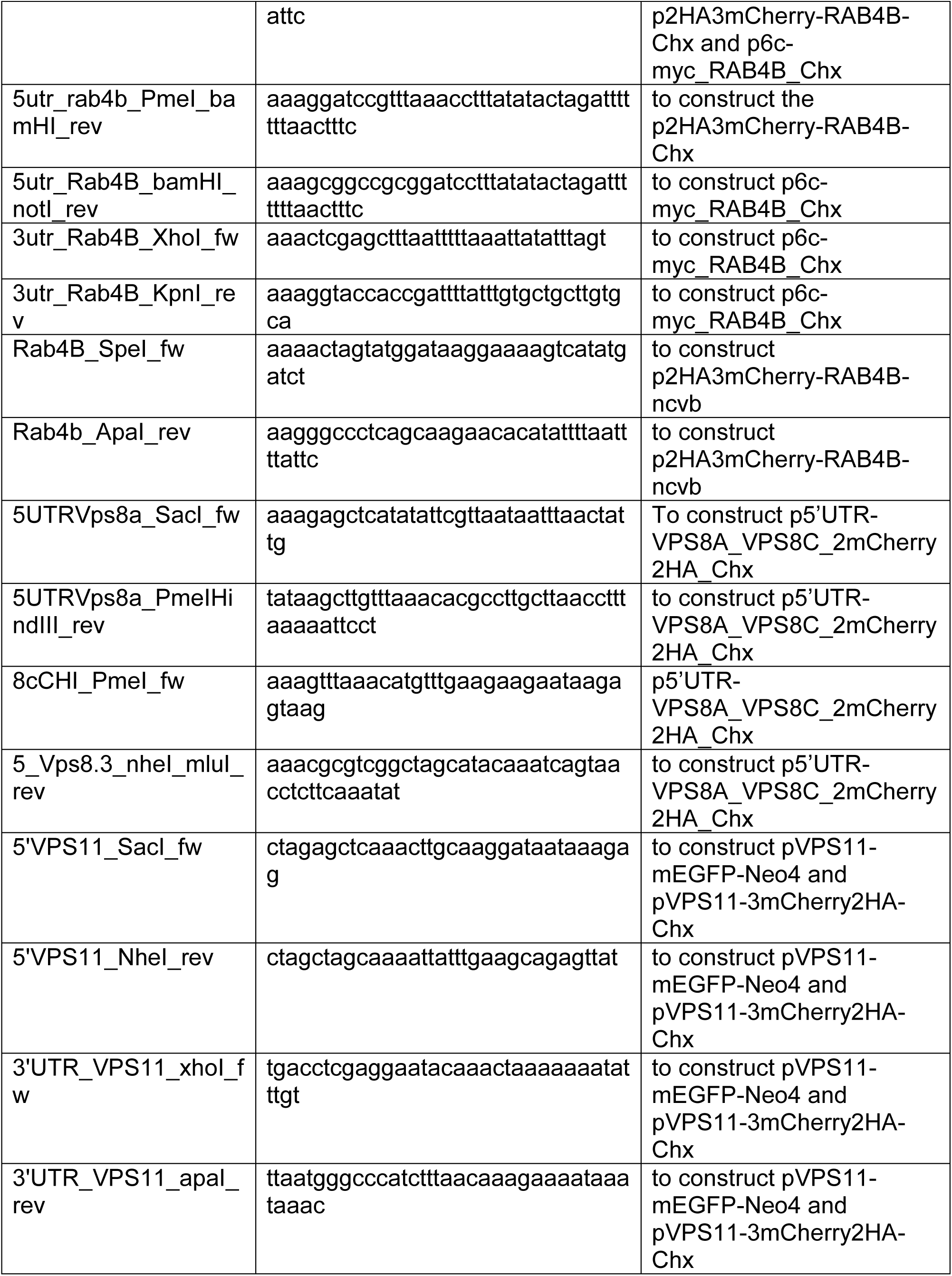
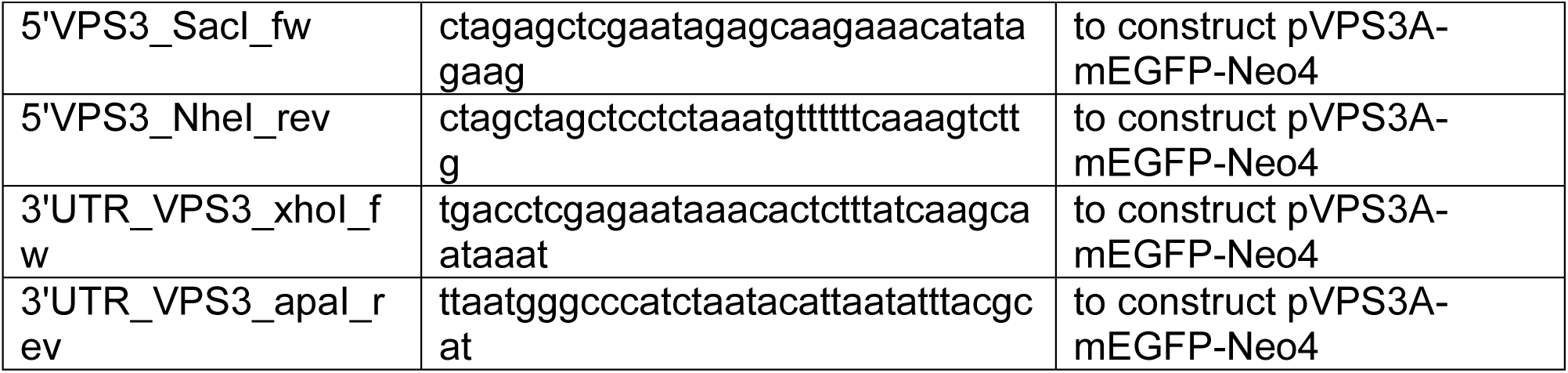
Primers used in this study

### Endogenous tagging of the Vps8 paralogs with FLAG epitope

The FLAG-ZZ tag, containing 3xFLAG, followed by the TEV (Tobacco Etch Virus cysteine protease) cleavage site and the IgG binding domain of protein A (ZZ-domain), was integrated at the C-termini of *VPS8B, VPS8C*, *VPS8D*, *VPS8E*, and *VPS8F* macronuclear ORFs by homologous recombination, using pVPS8B-FLAG-ZZ-Neo4, pVPS8C-FLAG-ZZ-Neo4, pVPS8D-FLAG-ZZ-Neo4, pVPS8E-FLAG-ZZ-Neo4 and pVPS8F-FLAG-ZZ-Neo4. The 5’ ends and the 3’ UTRs of *VPS8B, VPS8C*, and *VPS8D* were removed from pVPS8B-2mNeon-6myc-Neo4, pVPS8C-2mNeon-6myc-Neo4, pVPS8D-2mNeon-6myc-Neo4 vectors by using SacI/NheI and XhoI/ApaI, and cloned in the corresponding sites in pVPS11-FLAG-ZZ-Neo4, to replace the 5’ end and the 3’ UTR of *VPS11*, respectively. The same strategy was used to clone the 5’ ends and the 3’ UTRs of *VPS8E* and *VPS8F* at the SacI/MluI and EcoRV/XhoI sites in pVPS8A-FLAG-ZZ-Neo4 (Sparvoli et al., 2018) to replace the 5’ end and the 3’ UTR of *VPS8A*, respectively. The final constructs were digested with SacI and KpnI prior to biolistic transformation of CU428.1.

### Co-expression of c-myc-tagged Vps16a, Vps16b, Vps33a, Vps33b, Vps18a, Vps18b, Vps18c, or Vps18d with Vps8a-FLAG

The 6 c-myc tag was integrated at the C-termini of *VPS16A*, *VPS16B*, *VPS18A*, *VPS18B*, *VPS18C* and *VPS18D,* and at the N-termini of *VPS33A* and *VPS33B* by homologous recombination at the macronuclear ORFs, using pVPS16A-6c-myc-Chx, pVPS16B-6c-myc-Chx, pVPS18A-6c-myc-Chx, pVPS18B-6c-myc-Chx, pVPS18C-6c-myc-Chx, pVPS18D-6c-myc-Chx, p6c-myc-VPS33A-Chx and p6c-myc-VPS33B-Chx, respectively. PCR was used to amplify the 5’ ends (730-794bp minus the stop codon) and the 3’ UTRs (576-794bp) of *VPS16A*, *VPS16B*, *VPS18A*, *VPS18B*, *VPS18C*, and *VPS18D*. The 5’ ends and the 3’ UTR amplicons were cloned in the p6c-myc-Chx vector (Sparvoli et al., 2018) by Quick Ligation (New England, Biolabs Inc.) at SacI/NheI and XhoI/ApaI sites, respectively. *VPS33A* and *VPS33B* macronuclear ORFs were PCR-amplified and inserted by In-Fusion cloning (Clontech, Mountain View, CA) in the linearized p6c-myc-Chx vector at the SpeI site. PCR was used to amplify the 670-792bp of 5’ and 3’ UTRs of *VPS33A* and *VPS33B*. The 5’ and 3’ UTRs amplicons were then cloned in the corresponding p6c-myc-Chx vector containing the appropriate *VPS33* gene, by Quick Ligation at SacI-XbaI/BamHI and XhoI/ApaI sites, respectively. The final vectors pVPS16A-6c-myc-Chx, pVPS16B-6c-myc-Chx, pVPS18A-6c-myc-Chx, pVPS18B-6c-myc-Chx, pVPS18C-6c-myc-Chx, pVPS18D-6c-myc-Chx were digested with SacI and KpnI, p6c-myc-VPS33A-Chx with SacI and ApaI, p6c-myc-VPS33B-Chx with XbaI and KpnI, prior to biolistic transformation of Vps8a-FLAG-ZZ expressing cells. All primers are listed in Table 3.

### Endogenous tagging of Vps8c and Vps8e with mCherry in Vps8a-mNeon and Vps8c-mNeon expressing cells

3mCherry2HA tag was integrated at the C-termini of *VPS8C* and *VPS8E* macronuclear ORFs by homologous recombination using pVPS8C-3mCherry2HA-Chx and pVPS8E-3mCherry2HA-Chx. To construct the pVPS8C-3mCherry2HA-Chx, the 3FLAG-ZZ-Neo4 fragment in pVPS8C-FLAG-ZZ-Neo4 was replaced with 4491bp of the 3mCherry2HA-Chx fragment digested with NheI and XhoI from pVPS11-3mCherry2HA-Chx (see below), and cloned at the corresponding sites in pVPS8C-FLAG-ZZ-Neo4 by Quick Ligation (New England, Biolabs Inc.) pVPS8E-3mCherry2HA-Chx was obtained by first cloning the 2742bp of the 3mCherry2HA-3’UTR-BTU1 fragment from pVPS8C-3mCherry2HA-Chx at the NheI and PstI sites in pVPS8E-FLAG-ZZ-Neo4, and then by replacing the Neo4 drug resistance cassette with the PCR-amplified Chx cassette at the PstI and EcoRV sites via Quick Ligation. The final constructs were then linearized with SacI and KpnI, and pVPS8C-3mCherry2HA-Chx was biolistically transfected into Vps8ap-mNeon-expressing cells, while linearized pVPS8E-3mCherry2HA-Chx was transfected into Vps8c-mNeon and Vps8a-mNeon expressing cells. The primers are listed in Table 3.

### Expression of mCherry-tagged Vps8c at the VPS8A locus in Vps8c-mNeon expressing cells

A second 2mCherry2HA-tagged copy of *VPS8C* was introduced in the *VPS8A* locus in Vps8c-mNeon expressing cells by homologous recombination using p5’UTR-VPS8A-VPS8C-2mCherry2HA-Chx. To create p5’UTR-VPS8A-VPS8C-2mCherry2HA-Chx, we used p5’UTR-VPS8A-VPS8C-FLAG-ZZ-Chx as starting vector. Briefly, for the construction of p5’UTR-VPS8A-VPS8C-FLAG-ZZ-Chx we first replaced the Neo4 drug resistance cassette in pVPS8A-FLAG-ZZ-Neo4 (Sparvoli et al., 2018) with Chx at the SpeI and EcoRV sites, via digestion and Quick Ligation, to generate the pVPS8A-FLAG-ZZ-Chx vector. A 5218bp-long gene block was PCR amplified including the *VPS8C* macronuclear ORF together with the 813bp-long 5’UTR of VPS8A. The latter was digested with SacI and HindIII and cloned in pVPS8A-FLAG-ZZ-Chx at the corresponding sites, to integrate the additional copy of the *VPS8C* gene into the *VPS8A* genomic locus by homologous recombination. The reverse primer for the 5’UTR-VPS8A cloning contained a PmeI site upstream the HindIII site, thus the *VPS8C* gene ORF was cloned between PmeI and MluI sites of the corresponding vector by Quick Ligation. We then replaced the FLAG-ZZ-3’UTR-BTU1 fragment in p5’UTR-VPS8A-VPS8C-FLAG-ZZ-Chx with 2mCherry2HA-3’UTR-BTU1, obtained by digesting pGRL3-2mCherry2HA-Chx vector with BamHI and XmaI. The fragment was cloned by Quick Ligation into the BamHI/XmaI-linearized p5’UTR-VPS8A-VPS8C-FLAG-ZZ-Chx vector. The final construct p5’UTR-VPS8A-VPS8C-2mCherry2HA-Chx was linearized with SacI and KpnI prior to biolistically transforming *Tetrahymena*. Primers are listed in Table 3.

### Expression of GFP-tagged Vps3a and Vps11

Monomeric enhanced GFP (mEGFP) was integrated at the C-termini of *VPS3A* and *VPS11* macronuclear ORFs via homologous recombination using pVPS3A-mEGFP-Neo4 and pVPS11-mEGFP-Neo4, respectively. PCR was used to amplify the 5’ ends (666-778bp minus the stop codon) and the 3’ UTRs (785-690bp) of *VPS3A* and *VPS11*. The 5’ end and the 3’ UTR amplicons were cloned in the pmEGFP-Neo4 vector (Briguglio et al., 2013) by Quick Ligation (New England, Biolabs Inc.) at SacI/NheI and XhoI/ApaI sites, respectively. The final vectors pVPS3A-mEGFP-Neo4 and pVPS11-mEGFP-Neo4 were digested with SacI and KpnI prior to biolistic transformation of CU428.1.

### Expression of mCherry-tagged Vps11 or Vps8c in cells expressing Vps3a-GFP

Vps11-mCherry and Vps8c-mCherry were integrated at the corresponding endogenous loci in cells expressing Vps3a-GFP by homologous recombination, using pVPS11-3mCherry2HA-Chx and the previously mentioned pVPS8C-3mCherry2HA-Chx vectors. 3mCherry2HA was integrated at the C-terminus of the *VPS11* macronuclear ORF via homologous recombination using the pVPS11-3mCherry2HA-Chx vector. The vector was constructed by cloning the PCR-amplified 5’ end (778bp minus the stop codon) and 3’ UTR (690bp) of *VPS11* into the p3mCherry2HA-Neo4 vector (Sparvoli et al., 2018), at SacI/NheI and XhoI/ApaI sites, respectively. The resulting pVPS11-3mCherry2HA-Neo4 vector was then digested with PstI and XhoI to replace the Neo4 resistance cassette with the Chx cassette. The final vectors pVPS8C-3mCherry2HA-Chx and pVPS11-3mCherry2HA-Chx were linearized with SacI and KpnI prior to biolistic transformation of Vps3a-GFP expressing cells.

### Co-Expression of mCherry-Rab4b with Vps8a-mNeon and Vps8c-mNeon

The 2HA3mCherry tag was integrated at the N-terminus of the *RAB4B* (TTHERM_01097960) macronuclear ORF via homologous recombination using p2HA3mCherry-RAB4B-Chx. Constructs p6c-myc-RAB4B-Chx and p2HA3mCherry-RAB4B-ncvb were used as templates to construct the final RAB4B vector. First, the 5’UTR of *RAB4B* was PCR-amplified, digested with SacI and BamHI, and subsequently cloned by Quick Ligation (New England, Biolabs Inc.) at the corresponding sites in p6c-myc-RAB4B-Chx, to replace the 5’UTR and thereby introduce a PmeI site upstream of the BamHI site. *RAB4B* has a PmeI site within the genomic sequence, and this was used to linearize p6c-myc-RAB4B-Chx containing the new 5’UTR, and thus to introduce the 2767bp 2HA3mCherry-N-terminal RAB4B fragment. The latter was obtained by digesting the p2HA3mCherry-RAB4B-ncvb vector with PmeI. The correct orientation of the fragment was tested using MfeI and SpeI. The p2HA3mCherry-RAB4B-Chx final vector was linearized with SacI and KpnI prior to biolistic transformation of Vps8a-mNeon and Vps8c-mNeon expressing cells. The primers are listed in Table 3.

### Co-expression of mCherry-Rab7 or mCherry-Rab22a with Vps8c-mNeon

mCherry-Rab7 and mCherry-Rab22a were integrated at the metallothionein (*MTT1*) genomic locus in cells expressing Vps8c-mNeon by homologous recombination, using the previously described 2HA-3mCherry-RAB7-ncvb and 2HA-3mCherry-RAB22a-ncvb vectors (Sparvoli et al., 2018). The constructs were linearized with SfiI prior to biolistic transformation.

### Biolistic transformation

*Tetrahymena* transformants were generated and selected after biolistic transformation as previously described(Kaur et al., 2017; Sparvoli et al., 2018). Transformants were serially transferred 6x/week in increasing concentrations of drug and decreasing concentrations of CdCl_2_ (up to 2 mg/ml of paromomycin and 0.1 μg/ml CdCl_2_; up to 18-21 μg/ml of cycloheximide and 1 μg/ml CdCl_2_; up to 90 μg/ml of blasticidin and 0.1 μg/ml CdCl_2_) for at least 5 weeks before further testing. Successful integration and replacement of all endogenous alleles at each genomic locus was tested by RT-PCR as previously described (Sparvoli et al., 2018). At least three independent transformants were tested for each line.

### Co-immunoprecipitations

Vps8a-FLAG-ZZ was co-immunoprecipitated with Vps16b-myc, Vps33b-myc or Vps18d-myc, from detergent lysates of cells co-expressing Vps8a-FLAG-ZZ with either Vps16a-myc, Vps16b-myc, myc-Vps33a, myc-Vps33b, Vps18a-myc, Vps18b-myc, Vps18c-myc or Vps18d-myc, using anti-FLAG beads (EZ view Red Anti-FLAG M2 affinity Gel, Sigma) and anti-c-myc beads (Pierce Anti-c-Myc Agarose, Thermo Scientific), respectively, as previously described (Sparvoli et al., 2018). In brief, 300-500ml cultures were grown overnight to 2-3×10^5^ cells/ml. The cells were washed once with 10mM Tris-HCl, pH 7.4, pelleted and resuspended in cold lysis buffer (20 mM Tris-HCl pH 7.4, 50 mM NaCl, 1 mM MgCl_2_, 1 mM DTT, 1 mM EGTA, 0.2% NP-40, 10% glycerol, 4% BSA), supplemented with protease inhibitor cocktail tablets (Roche), and gently mixed for 45 min on ice. Lysates were cleared by centrifugation at 35000 rpm (Beckman Instruments type 45 Ti rotor) for 1.5 h at 4°C, split in two 50ml-falcon tubes, and separately mixed with 75 μl anti-FLAG and 400 μl anti-c-myc beads, pre-incubated with cold lysis buffer for 2 h at 4°C, respectively. The beads were then washed five times with 20 mM Tris-HCl pH 7.4, 1 mM EDTA, 500 mM NaCl, 0.1% NP-40, 1 mM DTT, 10% glycerol, and resuspended in 80 μl 100°C 2X LDS (lithium dodecyl sulfate) sample buffer containing 40 mM DTT.

### Immunoprecipitations from cryomilled cell powders

*Tetrahymena* were grown overnight to 2-3×10^5^ cells/ml, washed once with 10mM Tris-HCl pH 7.4 and re-pelleted. Supernatants were rapidly aspirated to leave a dense cell slurry. The slurries were transferred, drop-wise, into liquid nitrogen and milled to powders using a Cryogenic Grinder 6875 Freezer Mill. The cryopowders were resuspended in buffer B4 (20 mM Hepes pH 7.4, 250 mM NaCitrate, 1 mM MgCl_2_, 0.1% CHAPS, 1 mM DTT), supplemented with protease inhibitor cocktail tablets (Roche), gently mixed for 1h at 4°C, and then on ice until no solid matter was visible. Lysates were cleared by centrifugation at 35000 rpm (Beckman Instruments type 45 Ti rotor) for 1.5h at 4°C, and mixed with anti-FLAG beads (EZ view Red Anti-FLAG M2 affinity Gel, Sigma), pre-washed with cold lysis buffer, for 2 h at 4°C. The beads were then washed five times with 20 mM Tris-HCl pH 7.4, 1 mM EDTA, 500 mM NaCl, 0.1% NP-40, 1 mM DTT, 10% glycerol supplemented with protease inhibitor cocktail tablets (Roche). Samples destined for silver staining were washed one additional time with elution buffer (20 mM Hepes pH 7.4, 150 mM NaCl, 1.5 mM MgCl_2_, 0.1% CHAPS, 1 mM DTT, 5% glycerol, protease inhibitor tablets). Washed beads were then resuspended in elution buffer, or in 100°C LDS sample buffer containing 40 mM DTT, depending on the purpose of the experiment. For isolation of CORVET complexes for subsequent mass spectrometry, *Tetrahymena* from an overnight culture were inoculated in 10 L SPP, and grown to 2-3×10^5^ cells/ml for 24-26 h at 30°C with agitation at 75 rpm, and powders prepared as described above. 100-150 g of powder and 400 μl of anti-FLAG beads were used for each cell line. Proteins were eluted from the beads with 60μl 100°C LDS sample buffer containing 40 mM DTT, prior to SDS-PAGE and Coomassie staining. To isolate 8A-CC for subsequent analysis by glycerol gradient centrifugation, we used 30 g of cryopowder per experiment and 75 μl anti-FLAG beads. The complex was eluted by mixing the beads with 250 μl of 450 ng/μl 3XFLAG peptides in elution buffer for 2 h at 4°C. The elution step was repeated with additional 250 μl of 450 ng/μl 3XFLAG peptides, for 1h at 4°C. Roughly 450μl of these eluates were applied to glycerol gradients. For the isolation of CORVET complexes 8B, C, D, E and F-CC for SDS-PAGE and silver staining, we used 2 g of cryopowder per sample and 50 μl anti-FLAG beads. Complexes were eluted by mixing the beads with 150 μl of 450 ng/μl 3XFLAG peptides in elution buffer, for 2h at 4°C. 100°C LDS sample buffer containing 40 mM DTT was added to 150μl eluates prior to SDS-PAGE.

### Immunoprecipitation from cell pellets

This protocol was used to immunoprecipitate FLAG- and mNeon-tagged Vps8 paralogs, Vps8a-GFP, Vps8c-2mCherry, Vps8c-3mCherry and Vps8e-3mCherry, prior to visualization by western blotting. Wild-type cells were processed in parallel as control. 50-100 ml cell cultures were grown overnight to 3×10^5^ cells/ml, except the 3mCherry-tagged Vps8c and Vps8e that were grown to ∼7×10^5^ cells/ml. Cells were washed once in 10 mM Tris-HCl pH 7.4, pelleted, resuspended in buffer B4 (20 mM Hepes pH 7.4, 250 mM NaCitrate, 1 mM MgCl_2_, 0.1% CHAPS, 1 mM DTT) supplemented with protease inhibitor cocktail tablets (Roche), and gently mixed for 1h on ice. The lysates were cleared by centrifugation at 55000 rpm (Beckman Instruments TLA-100.4 rotor) for 1.5 h at 4°C, and mixed with 20μl antibody-conjugated beads, pre-washed with cold buffer B4, for 2 h at 4°C. Reagents used were: anti-FLAG beads (EZ view Red Anti-FLAG M2 affinity Gel, Sigma), anti-c-myc beads (Pierce Anti-c-Myc magnetic beads, Thermo Scientific), and anti-HA beads (EZ view Red Anti-HA Affinity Gel, Sigma), for FLAG-, mNeon-, and mCherry-tagged fusion proteins, respectively. To immunoprecipitate Vps8a-GFP, we used 120 μl anti-GFP beads (GFP-nAb agarose, Allele Biotechnology). Beads were washed five times with 20 mM Tris-HCl pH 7.4, 1 mM EDTA, 500 mM NaCl, 0.1% NP-40, 1 mM DTT, 10% glycerol, and resuspended in 50 μl 100°C LDS sample buffer containing 40 mM DTT.

### Glycerol gradient centrifugation

Continuous 11-ml glycerol gradients were made by layering 10%, 20%, 30% 40% (v/v) glycerol solutions in 14×89 mm tubes (Beckman Instruments). The glycerol solutions were prepared in 20 mM Hepes pH 7.4, 150 mM NaCl, 1.5 mM MgCl_2_, 0.1% CHAPS, 1 mM DTT. The tubes were gently laid on their side for 1.5 h, and then stood upright for overnight at 4°C. Protein samples were overlaid on the gradients and the tubes centrifuged for 18 h at 37000 rpm using a SW 41 Ti rotor (Beckman Instruments), at 4°C. 250-μl fractions were harvested and analyzed by western blotting and silver staining.

### Western blotting

Protein samples were analyzed by western blot as previously described (Sparvoli et al., 2018). Mouse mAb anti-GFP (BioLegend), mouse mAb anti-c-myc (9E10, Sigma), rabbit anti-FLAG (Sigma), and mouse mAb anti-HA (HA.11, BioLegend) antibodies, were diluted 1:5000, 1:5000, 1:2000, 1:2000, respectively, in blocking solution. Proteins were visualized with either anti-rabbit IgG (whole molecule)-HPeroxidase (Sigma) or ECL Horseradish Peroxidase-linked anti-mouse (NA931) (GE Healthcare Life Sciences, Little Chalfont, UK) secondary antibody diluted 1:20000, and SuperSignal West Femto Maximum Sensitivity Substrate (Thermo Scientific).

### Silver staining

Proteins samples were loaded on 8% Tris-Glycine gels (Invitrogen) and stained with Pierce Silver Stain for Mass Spectrometry (Thermo Scientific), according to the manufacturer’s instructions.

### Mass spectrometry

Protein samples were loaded on a 4-20% gel for SDS-PAGE, allowed to migrate for ∼1 cm into the gel, and briefly stained with Coomassie Blue R-250 solution (0.1% w/v Coomassie, 10% acetic acid, 50% methanol). A single 1 cm gel slice per lane was excised from the Coomassie stained gel, destained, and then subjected to tryptic digest and reductive alkylation. Liquid chromatography tandem mass spectrometry (LC-MS/MS) was performed by the Proteomic Facility at the University of Dundee. The samples were subjected to LC-MS/MS on a Ultimate3000 nano rapid separation LC system (Dionex) coupled to a LTQ Velos mass spectrometer (Thermo Fisher Scientific). Mass spectra were processed using the intensity-based label-free quantification (LFQ) method of MaxQuant version 1.6.1.0 (Cox et al., 2014; Cox and Mann, 2008) searching the *T*. *thermophila* annotated protein database from ciliate.org (Eisen et al., 2006; Stover et al., 2006). Minimum peptide length was set at six amino acids, isoleucine and leucine were considered indistinguishable and false discovery rates (FDR) of 0.01 were calculated at the levels of peptides, proteins and modification sites based on the number of hits against the reversed sequence database. If the identified peptide sequence set of one protein contained the peptide set of another protein, these two proteins were assigned to the same protein group. Perseus (Tyanova et al., 2016) was used to calculate *P* values applying *t*-test based statistics and to draw statistical plots. Proteomics data have been deposited to the ProteomeXchange Consortium via the PRIDE partner repository (Vizcaino et al., 2016) with the dataset identifier PXD014895.

### Feeding with DsRed-bacteria

*E. coli* expressing DsRed-express2 (Strack et al., 2008) (a gift from B. Glick, The University of Chicago, Chicago, IL) were grown in 15 ml LB broth, supplemented with ampicillin, and induced overnight at 37°C with 1mM IPTG. *Tetrahymen*a expressing Vps8b-mNeon or Vps8e-mNeon were grown in 20 ml SPP overnight to 2.5×10^5^ cells/ml, and were incubated with 1% dsRed-expressing *E. coli* for 30 minutes at 30°C, prior to fixation.

### Fixed cell imaging

Cells (3×10^5^) endogenously expressing Vps11-GFP, Vps3a-GFP alone or with Vps11-mCherry or Vps8c-mCherry, mNeon-tagged Vps8 paralogs alone or in combination with mCherry-tagged Vps8c, Vps8e and Rab4b, were washed once with 10 mM Tris-HCl pH 7.4, and fixed with ice-cold 4% paraformaldehyde in 1X PBS for 30 min. Cells fed with dsRed-*E.coli* were collected by centrifugation, washed once with SPP and fixed. For the simultaneous localization of Vps8c-mNeon with either mCherry-Rab22a or mCherry-Rab7, the expression of the Rab-GTPases was induced by incubating the cells with 1 μg/ml CdCl_2_ for 2 h in SPP, prior to fixation. The visualization of fusion proteins was not enhanced with immunolabeling. Cells were mounted with Trolox (1:1000) to inhibit bleaching and imaged at room temperature on a Marianas Yokogawa type spinning disk inverted confocal microscope, 100X oil with NA=1.45, equipped with two photometrics Evolve back-thinned EM-CCD cameras, with Slidebook6 software (Zeiss, Intelligent Imaging Innovations, Denver, CO). Z stack images and z projection images were denoised, adjusted in brightness/contrast and colored with the program Fiji (Schindelin et al., 2012).

### Live cell imaging

*Tetrahymena* expressing the mNeon-tagged Vps8 paralogs, or co-expressing Vps8c-mNeon and mCherry-Rab7, were grown overnight to 1-2×10^5^ cells/ml and transferred to S medium for 2 h prior to imaging. The expression of Rab7 was induced by adding 1 μg/ml CdCl_2_ to the S medium. Cells were immobilized in thin 3% low melting agarose gel pads, as described previously (Kaur et al., 2017), and imaged within 15 minutes. Z stack images (12 stacks along the z axis at 0.5 μm intervals) and time-lapse videos (30 frames at 1.24 sec interval for mNeon-tagged Vps8 paralogs; 200 frames at 0.17 sec/interval for Vps8c-mNeon/mCherry-Rab7) were collected at room temperature with a Marianas Yokogawa type spinning disk inverted confocal microscope, 100X oil with NA=1.45, equipped with two photometrics Evolve back-thinned EM-CCD cameras, with Slidebook6 software (Zeiss, Intelligent Imaging Innovations, Denver, CO). Images and movies were denoised, and adjusted in brightness/contrast with the program Fiji (Schindelin et al., 2012). Images shown are single slices/frames for clarity. Videos of Vps8c-mNeon/mCherry-Rab7-expressing cells were created by simultaneously recording in two fluorescent channels, which were subsequently merged in a single multicolor movie using the program Fiji.

### Colocalization analysis

To estimate the extent of colocalization, we calculated the Mander’s coefficients M1 and M2 with the Fiji-JACoP plugin, as previously described (Sparvoli et al., 2018). 10-15 cells were analyzed to measure the extent of overlap between Vps8 paralogs and Rab-GTPases. In particular, for Vps8a/Rab4b, Vps8c/Rab4b, Vps8c/Rab22a, and Vps8c/Rab7, 158, 139, 268, and 232 non-overlapping images/sample were collected, respectively. 18-25 cells were imaged to calculate the extent of overlap between distinct Vps8 paralogs: for Vps8a/Vps8c, Vps8a/Vps8e, and Vps8c/Vps8e, coefficients M1 and M2 were derived from 399, 246, and 337 non-overlapping images/sample, respectively. To measure the colocalization between Vps8c-mNeon and Vps8c-mCherry, 14 cells were analyzed and the coefficients were derived from 261 non-overlapping images. 20 and 21 cells were used to calculate the overlap of Vps3a-GFP with either Vps11-mCherry or Vps8c-mCherry, respectively, and coefficients M1 and M2 were derived from 325 and 321 non-overlapping images/sample. Mander’s coefficients M1 and M2 are reported as Mean values.

### Particle counting

To estimate the number of fluorescent puncta in Vps8a-mNeon and Vps8a-GFP expressing cells, we used the Fiji tool “SpotCounter”, setting the tolerance noise to 1200-1500, and box size to 3. The plugin counts spots by detecting local maxima, which are accepted when the maximum is higher than a user-defined number (tolerance noise) over the average of the 4 corners of the box. The number of particles was calculated using maximum intensity z projections, generated from z stacks of 15 cells/sample, and it is reported as Mean value.

### Transcription profiles

Gene expression profiles were downloaded from the *Tetrahymena* Functional Genomics Database (TFGD, http://tfgd.ihb.ac.cn/) (Miao et al., 2009; Xiong et al., 2011b). For plotting the graphs, each profile was normalized by setting the gene’s maximum expression level to 1.

## Acknowledgements

At the University of Chicago, we thank Klaus Nielsen and Jon Staley for help with cryomilling, Matthew Sullivan and Alex Ruthenburg for help with FPLC, Aarthi Kuppannan for help with cell culture, and Vytas Bindokas for help with confocal microscopy at the Integrated Light Microscopy Core Facility. We thank Naomi Stover (Bradley University) for help with gene annotation, and Lev Tsypin (Caltech) for stimulating discussion. We thank the FingerPrints Proteomic Facility at the University of Dundee for excellent technical support. Work in APT’s laboratory was supported by NIH GM105783 and in MCF’s laboratory by Wellcome Trust Investigator Award 204697/Z/16/Z.

---

**Supplementary figure 1. Related to figure 2.**
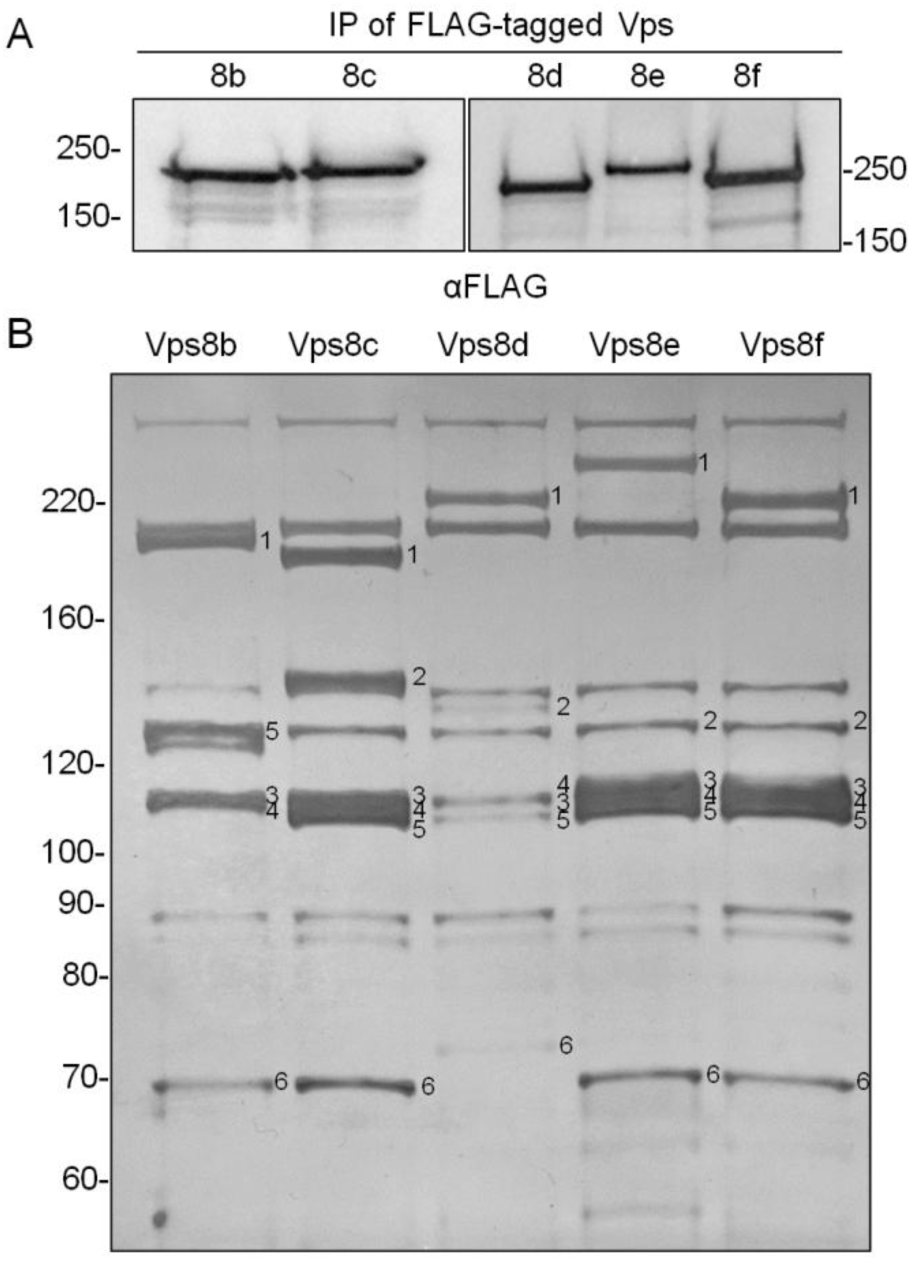
A) Immunoprecipitation of Vps8b, Vps8c, Vps8d, Vps8e, Vps8f. Cells were transformed to FLAG-tag the Vps8 paralogs at their endogenous loci. Detergent cell lysates were treated with anti-FLAG beads, and LDS-eluted proteins were analyzed by SDS-PAGE and Western blot with anti-FLAG antibodies. The expected full-length proteins were detected. B) FLAG-tagged Vps8b, Vps8c, Vps8d, Vps8e, Vps8f were immunoisolated from solubilized cryopowders using anti-FLAG beads. Bound proteins were eluted with 3XFLAG peptides, and 25% was loaded on 8% gels for SDS-PAGE followed by silver staining. For each lane, the numbers 1, 2, 3, 4, 5, and 6 indicate the Vps8, Vps18, Vps16, Vps11, Vps3, and Vps33 subunits for that complex, respectively. All six CORVET subunits were detected for Vps8c, Vps8d, Vps8e, and Vps8f. In the Vps8b-FLAG eluate, the Vps18 subunit (Vps18b) was not clearly detected at the expected size, but may migrate anomalously (note species migrating close to band 5, unique to this sample). For the Vps8d sample, Vps8d appears to be present in excess of all other subunits.

**Supplementary figure 2. Related to figure 3.**
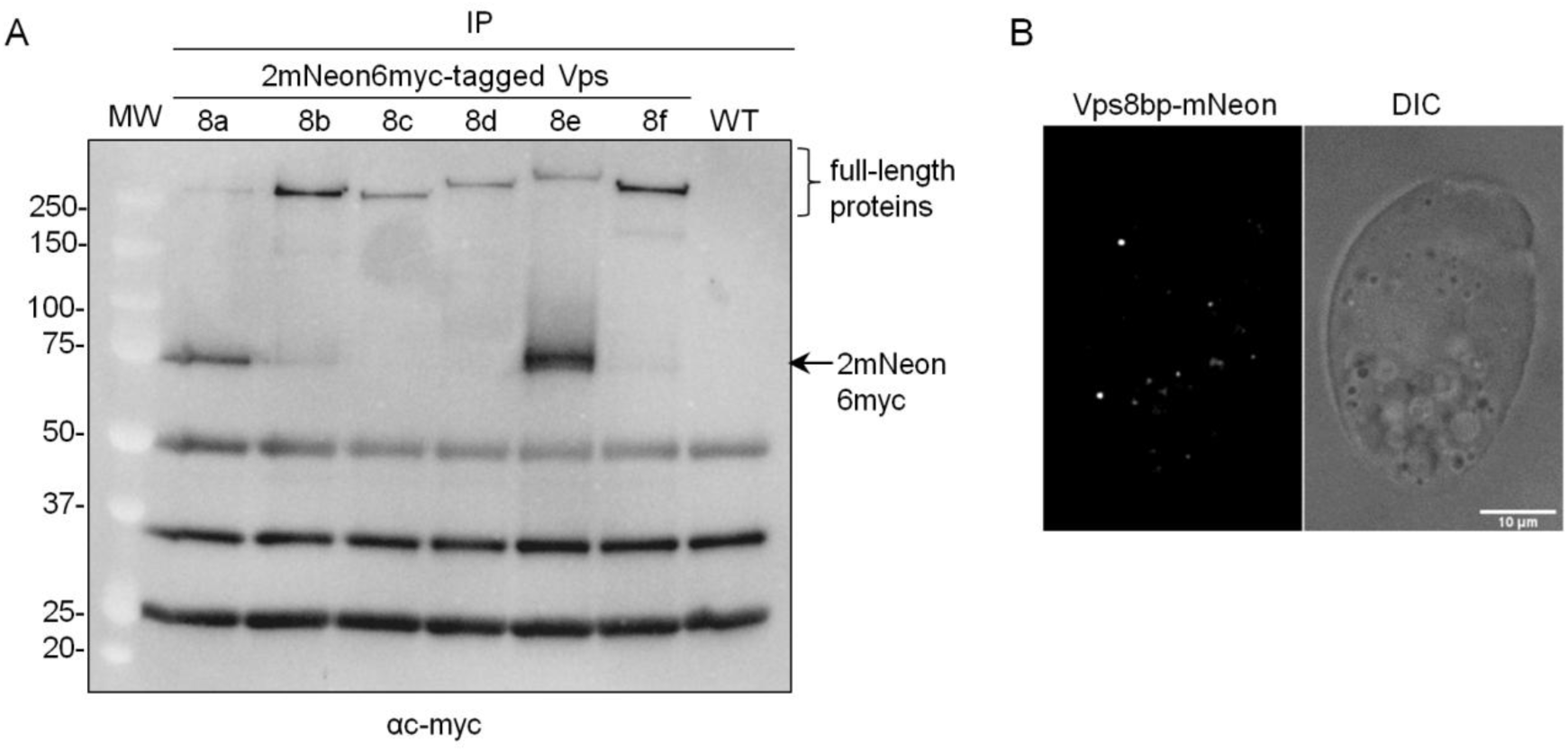
A) Immunoprecipitation of mNeon-tagged Vps8 paralogs. Detergent cell lysates were incubated with anti-c-myc beads, and bound proteins eluted with LDS-sample buffer. Samples were analyzed by SDS-PAGE and Western blotting with anti-c-myc antibodies. On the right, the black arrow indicates a band whose size corresponds to a cleaved 2mNeon6myc tag, visible in Vps8a and Vps8e samples, while a weaker band is present in Vps8b. B) Single frame from a time-lapse movie, with paired differential interference contrast (DIC) image, of a cell endogenously expressing mNeon-tagged Vps8b. Cells were incubated in S-medium for 2h prior to imaging. In favorable focal planes of live cells, Vps8b can be visualized in cytoplasmic puncta, which are more prominent in fixed cells (figure 1B).

**Supplementary figure 3. Related to figure 5.**
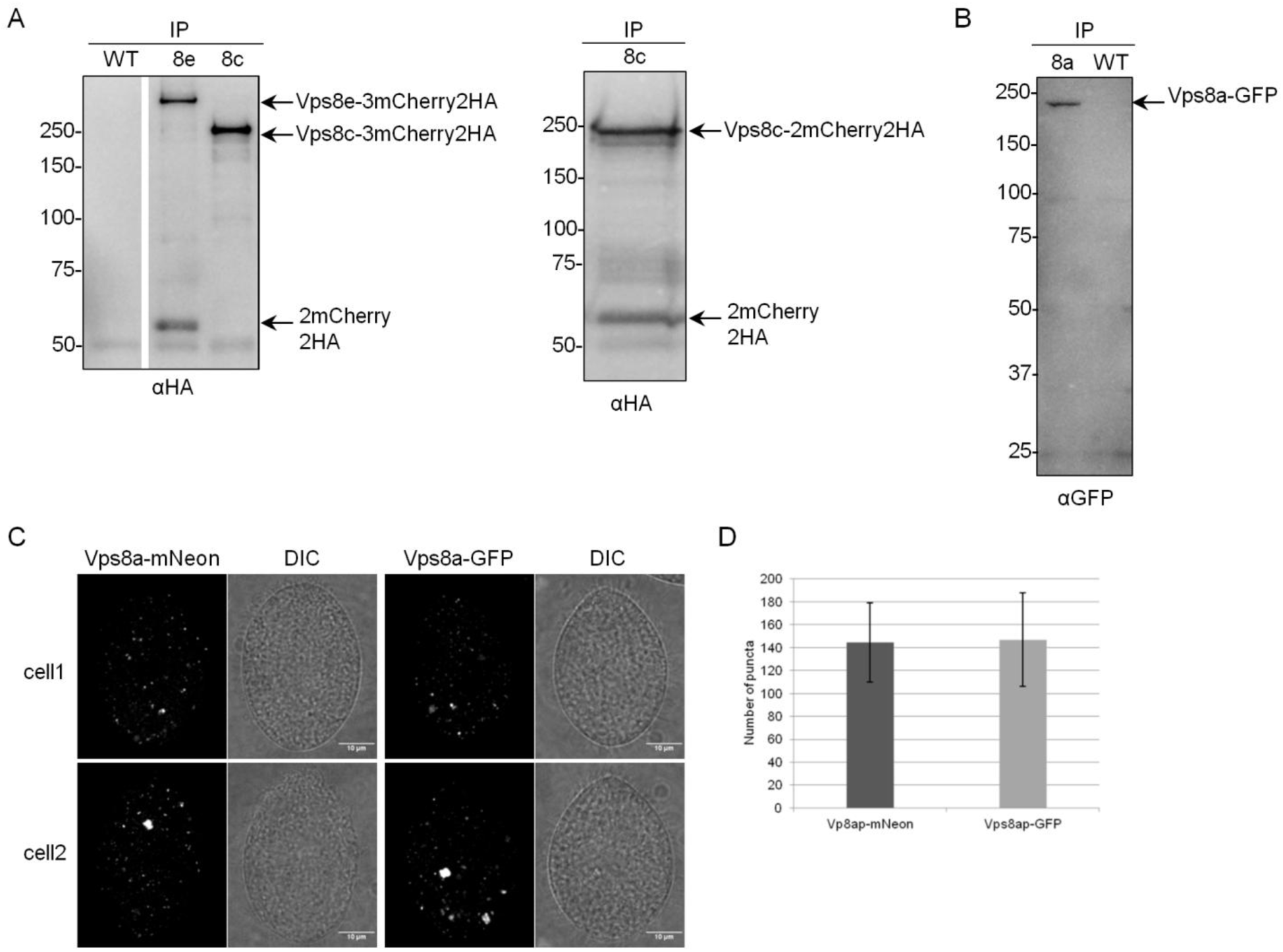
A) Immunoprecipitation of 3mCherry-tagged Vps8c and Vps8e (left), and 2mCherry-tagged Vps8c (right). Cells were transformed to co-express Vps8a-mNeon together with either Vps8e-3mCherry or Vps8c-3mCherry, or to express Vps8c-2mCherry at the *VPS8A* locus in Vps8c-mNeon-expressing cells. Detergent cell lysates were incubated with anti-HA beads, and bound proteins were eluted with LDS-sample buffer. Samples were subsequently analyzed by SDS-PAGE and Western blotting with anti-HA antibodies. In the 3mCherry-Vps8e sample, a band whose size corresponds to 2mCherry2HA is visible (western blot on the left), suggesting that cleavage occurred between the first and second copy of mCherry in the tag. An equivalent band is also visible in the western blot on the right. B) Immunoprecipitation of Vps8a-GFP expressed endogenously in SB281 cells. SB281 has a loss-of-function mutation in the *VPS8A* gene, and expression of the GFP-tagged copy rescues the mutant phenotype(Sparvoli et al., 2018). Detergent cell lysates were treated as in (B), and incubated with anti-GFP beads. Following SDS-PAGE and Western blotting with anti-GFP antibodies, only full length protein was detected. C) Maximum intensity projections of confocal z-stacks of fixed cells expressing mNeon-tagged Vps8a (left panel), and GFP-tagged Vps8a (right panel). The DIC images represent confocal cross sections for clarity. The fluorescent puncta for each fusion protein were counted, as summarized in (D). Scale bars, 10 μm. D) Quantification of fluorescent puncta/cell measured in 15 cells/sample, using the Fiji SpotCounter plugin. Error bars represent Standard Deviations. Despite the fact that the Vps8a-mNeon fusion undergoes some cleavage, the number of puncta labeled by this protein is not significantly different from uncleaved Vps8a-GFP, suggesting that the distribution of Vp8a-mNeon reflects the full cellular distribution of Vps8a.

**Supplementary figure 4. Related to figure 7.**
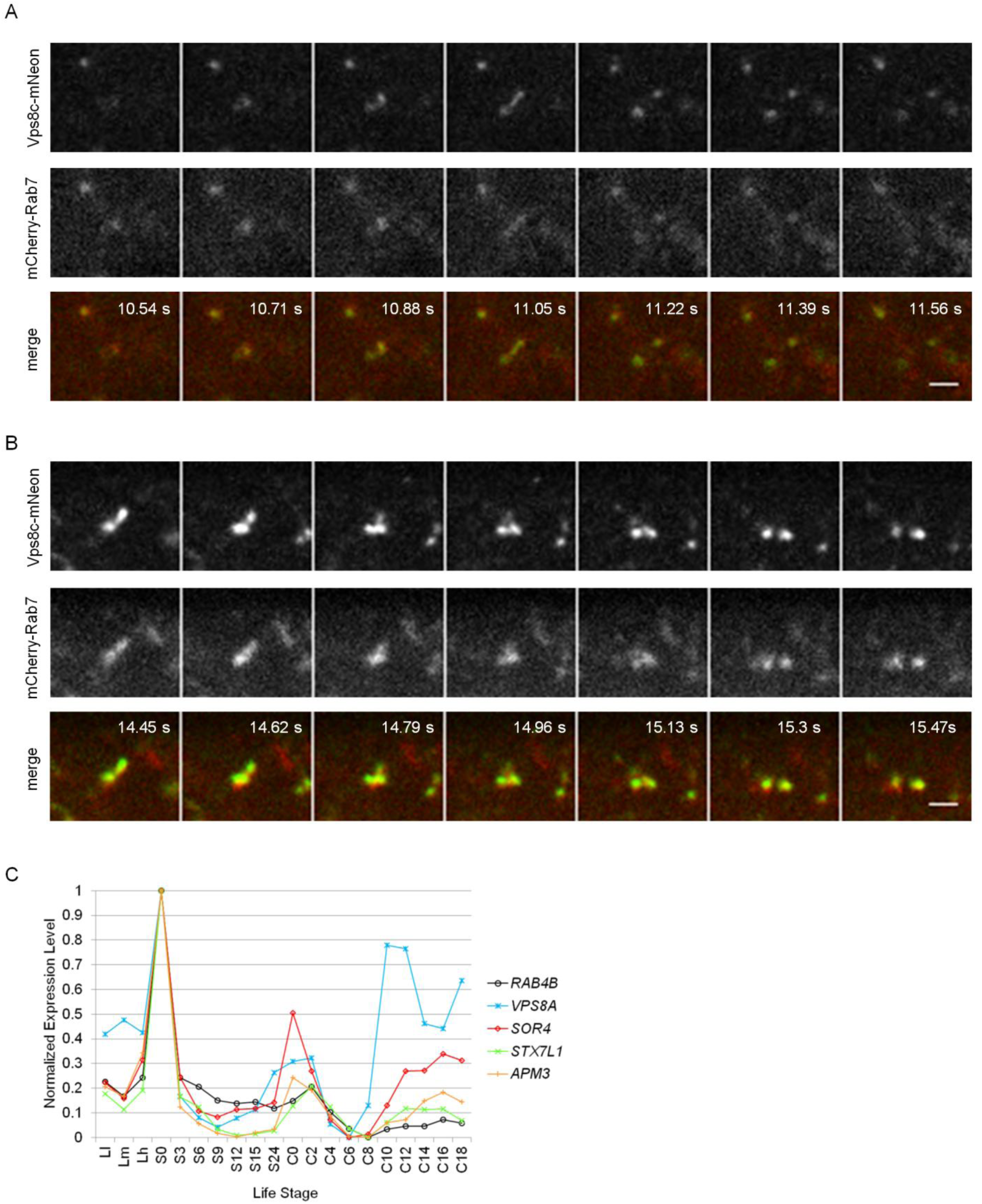
A) and B) Live images of selected areas of a cell co-expressing Vps8c-mNeon (top) and mCherry-Rab7 (middle)(bottom=merge). The two panels show events of Rab7/Vps8c-tubule outgrowth and apparent fission, occurring at different time points in different regions of the same cell. The sequential images for each fluorescent channel were simultaneously acquired at 0.17 sec/frame interval. The time intervals corresponding to the appearance and subsequent split of the tubulovesicular structures in the video, are indicated in white in each merged frame. Scale bar, 2 μm. C) Rab4b shares the transcriptional profile of mucocyst-associated genes. The expression profile of *RAB4B* (black line) is similar to that of mucocyst-related genes, here illustrated by *VPS8A*, *SOR4*, *STX7L1*, and *APM3*. Transcription profiles were downloaded from http://tfgd.ihb.ac.cn, based on sampling from growing cultures (low Ll, medium Lm, and high Lh culture density), starvation over 24 hr (S0–S24), and time points during conjugation (C0–C18). For clarity in plotting, each trace was normalized to that gene’s maximum expression level.

